# Light Potentials of Photosynthetic Energy Storage in the Field: What limits the ability to use or dissipate rapidly increased light energy?

**DOI:** 10.1101/2021.08.26.457798

**Authors:** Atsuko Kanazawa, Abhijnan Chattopadhyay, Sebastian Kuhlgert, Hainite Tuitupou, Tapabrata Maiti, David M. Kramer

## Abstract

The responses of plant photosynthesis to rapid fluctuations in environmental conditions are thought to be critical for efficient capture of light energy. Such responses are not well represented under laboratory conditions, but have also been difficult to probe in complex field environments. We demonstrate an open science approach to this problem that combines multifaceted measurements of photosynthesis and environmental conditions, and an unsupervised statistical clustering approach. In a selected set of data on mint (*Mentha* sp.), we show that the “light potential” for increasing linear electron flow (LEF) and nonphotochemical quenching (NPQ) upon rapid light increases are strongly suppressed in leaves previously exposed to low ambient PAR or low leaf temperatures, factors that can act both independently and cooperatively. Further analyses allowed us to test specific mechanisms. With decreasing leaf temperature or PAR, limitations to photosynthesis during high light fluctuations shifted from rapidly-induced NPQ to photosynthetic control (PCON) of electron flow at the cytochrome *b*_*6*_*f* complex. At low temperatures, high light induced lumen acidification, but did not induce NPQ, leading to accumulation of reduced electron transfer intermediates, a situation likely to induce photodamage, and represents a potential target for improving the efficiency and robustness of photosynthesis. Finally, we discuss the implications of the approach for open science efforts to understand and improve crop productivity.

## Introduction

While oxygenic photosynthesis supplies energy to drive essentially all biology in our ecosystem, it involves highly energetic intermediates that can generate highly toxic reactive oxygen species (ROS) that can damage the organisms it powers [1]. Thus, the energy input into photosynthesis must be tightly regulated by photoprotective mechanisms that act at several key steps in the light reactions. The balance and kinetics of this regulation is an active target for crop improvement.

One class of photoprotective processes, known as nonphotochemical quenching (NPQ), dissipates absorbed light energy as heat, thus diverting energy away from photosystem II (PSII) [2], decreasing the accumulation of reactive intermediates. This photoprotective capacity comes at the cost of decreased photochemical efficiency, and thus the organisms must regulate NPQ to balance the avoidance of photodamage with efficient energy conversion [3,4]. There are several forms of NPQ that differ in their mechanisms and rates of activation and deactivation. The most rapid NPQ form is qE, which is activated by acidification of the thylakoid lumen by the proton gradient (ΔpH) component of the thylakoid proton motive force (*pmf*) [2]. Lumen acidification activates the violaxanthin deepoxidase or VDE [5–8] resulting in the conversion of violaxanthin (Vx) to antheraxanthin (Ax) and zeaxanthin (Zx); and protonation of PsbS, an antenna-associated protein required for qE [2], which appear to act cooperatively in setting the extent of qE. The conversion of Vx to Ax and to Zx is typically much slower than the rapidly reversible protonation of PsbS [2], and during prolonged illumination, the responses of qE will likely be limited by the rate of acidification and de-acidification of the thylakoid lumen, which are, in turn, governed by ion movements in the chloroplasts [9–11]. Slower forms of NPQ have also been demonstrated [12], including qI, which is related to the photodamage and repair of photosystem II (PSII) or qZ, which related to the accumulation of Zx (independently from qE) [13], qH, related to cold and high light stress [13], and qT, related to antenna state transitions [14].

The acidification of the thylakoid lumen also controls electron transfer at the cytochrome *b*_*6*_*f* complex, a process called photosynthetic control (PCON) [15–20], which prevents the buildup of electrons on the acceptor side of photosystem I (PSI) that can lead to photodamage [15,21–23]. Interestingly, PCON and qE (both responses to lumen acidification) are expected to have opposing effects on Q_A_ redox state. High levels of PCON in the absence of qE would lead to accumulation of plastoquinol (PQH_2_) and the reduced form of the PSII electron acceptor, Q_A_-, which can potentiate photodamage. Thus, these two processes must be tightly coordinated, with qE being activated at lumen pH somewhat less acidic than PCON [15].

Plants in natural environments are exposed to rapidly changing environmental conditions, especially light which can change by orders of magnitude in less than a second. It has become clear that rapid and unpredictable fluctuations in light intensity can be more damaging than more gradual changes [22,24–31]. This sensitivity can partly be related to the buildup of reactive redox intermediates and thylakoid *pmf*, which can occur following low-to-high light transitions much more rapidly than the activation of photoprotective NPQ and PCON, leaving the photosynthetic apparatus prone to photodamage. Also, the slow recovery of NPQ following a decrease in light intensity can lead to substantial losses of photosynthetic efficiency [32]. Recently, it has been reported that engineering plants with increased expression levels of VDE and zeaxanthin epoxidase (ZE), resulted in accelerated formation and reversal of qE accompanied by increased plant productivity [3], suggesting that it may be possible to increase yield in crops by modifying photosynthetic regulatory responses.

On the other hand, we lack comprehensive surveys of the range of natural response of photosynthesis to real environmental fluctuations, in part because of a lack of deployable scientific equipment and methods to probe these processes in the field. Consequently, it has not been possible to assess the mechanistic bases of extant natural variations in these processes, their possible benefits or tradeoffs, or which of these may be most useful for crop improvement.

Here, we introduce a method and proof-of-concept field data results to address the following questions: Can we assess the extent of natural variations in rapid responses to fluctuations in photosynthetically-active radiation (PAR) intensity for both electron flow and photoprotection? How do these limitations depend on environmental conditions? What are the mechanisms that underlie these variations in responses to rapidly fluctuating light in the field?

Here, we introduce an approach to both measure and analyse these variations, focusing on one species, *Mentha* sp., under a limited set of conditions, and applied these to testing among a set of mechanisms that can be distinguished based on a range of optical measurements available using the MultispeQ 2.0 device, including: 1) PSI acceptor-side limitations to electron transfer; 2) Increased NPQ which limits the input of light energy into photosystem II (PSII); and 3) Photosynthetic control in which acidification of the lumen slows electron transfer at the level of plastoquinol (PQH_2_) oxidation by the cytochrome *b*_*6*_*f* complex.

The results show that the approach can effectively be used to assess the range of variations in ‘light potentials,’ the extent to which increased light leads to increased photosynthetic responses, under field conditions, as well as to test specific hypothetical models, setting up a broad-scale, multiple participant, open science approach to exploring the responses across multiple species, genotypes and environments. The results also reveal, at least in *Mentha*, unexpected leaf temperature-dependent limitations in the rapid formation of NPQ that result in the accumulation of reduced PSII electron acceptor, Q_A_ and a high thylakoid *pmf*, conditions likely to promote the formation of reactive oxygen species.

## Materials and Methods

### Plants and leaf sampling

Measurements were made in a population of *Mentha* sp. (mint) plants that have been maintained at the MSU Horticulture Gardens for at least 10 years. The GPS locations of all measurements are included in the online data set (https://photosynq.org/projects/rapid-ps-responses-pam-ecst-npqt-mint-dmk). Although it was not practical to exhaustively capture the lifecycle of the plants, the experimental strategy sampled a sufficiently wide range of conditions to allow clear patterns emerged in the relationships between phenotypes and environmental parameters, as described below. The experiment took place over a nine-day experimental window (Figure S1A), sampling a range of times of day, temperatures etc. (Fig. S1B). Measurements were made at multiple, alternating canopy levels and positions (subjectively at high, middle and low canopy levels) from early morning, though later afternoon (Fig. S1B), and at multiple locations across the plots on each day.

### Measurements of photosynthetic and related parameters using MultispeQ 2.0

Optical measurements were made using MultispeQ V2.0 hand-held instruments (https://photosynq.com), based on that presented by Kuhlgert et al [33] and calibrated using the CaliQ calibration system (https://photosynq.com/caliq). The Light Potential (LP) protocol used in the experiments can be found in the online project information (rapid-ps-responses-with-ecs-fast-ecs-dirk-and-npqt-dmk) as illustrated in Figure 1. The protocol was designed to strike a balance between needs for sampling large numbers of leaves, the desire for detailed spectroscopic measurements and the length of time the plant could be exposed to increased or decreased PAR. The full protocol, with measurements at ambient, after 10 s full sunlight and 10 s dark required about 35-40 s, at the limit of the time scale over which most researchers could steadily clamp a leaf in the instrument. The implications of the 10 s illumination and recovery time are discussed in the Results and Discussion sections.

**Figure 1.**
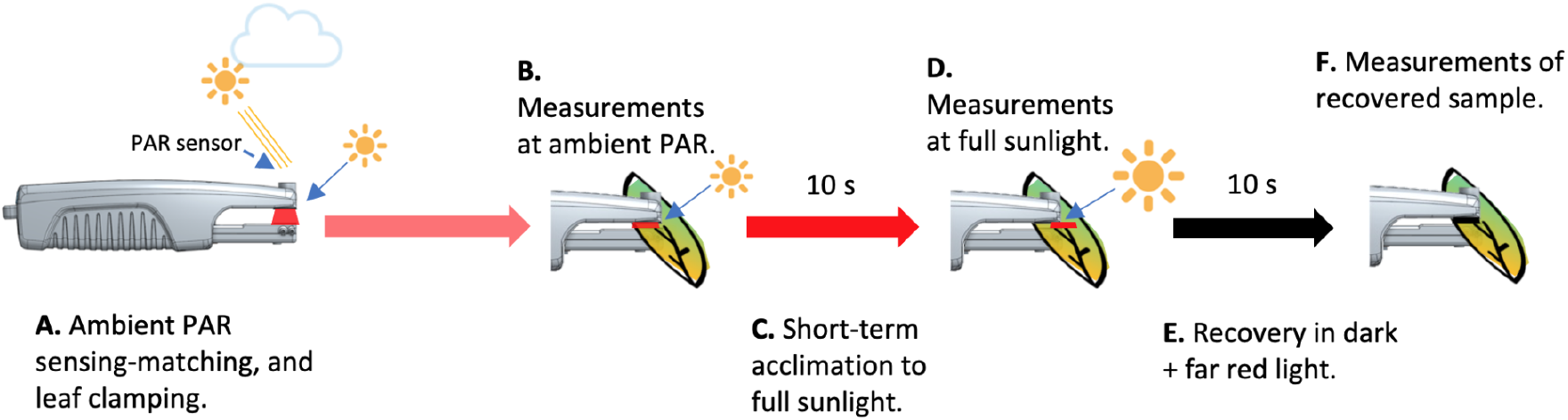
Experimental procedure of NPQ Light Potential designed to detect the change in NPQ induced under different light intensities. A) A sensor on the MultispeQ continuously detects the ambient photosynthetically active radiation (PAR) in the field and reproduces this PAR value using internal LED. B)When the leaf clamp is closed over a leaf, the experiment begins by recording the local PAR, leaf temperature, ambient temperature, leaf angle and GPS position. After a short period of illumination at the measured ambient PAR, the first set of optical measurement are recorded. C) Once completed, the leaf is exposed to a period of high PAR (2,000 μmol·m^−2^·s^−1^, equivalent to full sunlight) for 10 s. D) The optical measurements are repeated at high PAR. E) The leaf is then dark adapted (actinic light is switched off), with weak far red background light for 10 s (Panel E). F) A final set of optical measurements is made to assess rapid dissipation of NPQ and reoxidation of accumulated reduced intermediates. Each set of optical measurements includes chlorophyll fluorescence and absorbance changes to give estimates of ϕ_II_, LEF, NPQt, q_L_ (Q_A_ redox state); ECSt, P_700_ redox state, and g_H+_ (relative proton conductivity of the thylakoid ATP synthase), as described in Materials and Methods. Measurements taken at ambient and light are designated as “measurement (ambient), LEF (ambient), NPQt (ambient), q_L_ (ambient).

In the first stage of the protocol (Fig. 1A), the MultispeQ was programmed to continuously (at about 5 Hz) measure photosynthetically-active radiation (PAR) and reproduced these levels using a red actinic LED (655 nm emission peak) illuminating the adaxial surface of the leaf. When the MultispeQ detected that a leaf was clamped in the chamber, a series of measurement sequences were initiated. After a few seconds of illumination at ambient PAR (PAR_amb_) to allow for settling and setting of gains, the first set of measurements was made, estimating at PAR_amb_ LEF (LEF_amb_), NPQt (NPQ_amb_) and other photosynthetic parameters (Fig. 1B).

The actinic light was then increased to approximately full sunlight (2,000 µmol·m^-2^·s^-1^ red light) for 10 s (Fig. 1 C), after which the photosynthetic measurements were repeated (Fig 1D), yielding measurements of LEF_high_, NPQ_high_ etc. We chose full sunlight, rather than an artificially intense super-saturation light, to estimate light potentials that could occur in the field, and not the absolute maximum, and to avoid non-physiological or photoinhibitory effects. Thus, the light potentials of various processes will be limited as PAR_amb_ approaches full sunlight.

In the third stage of the experiment, the actinic light was then switched off, and a weak far-red light switched on for 10s (Fig. 1E), following another repetition of the measurements to assess the extent of NPQ_t_ after relaxation (NPQ_rec_, Fig. 1F). Environmental parameters including PAR, temperature, humidity, leaf temperature, leaf angle and GPS location were measured either prior to or following the physiological measurements.

Chlorophyll fluorescence changes were measured using MultispeQ 2.0 devices to estimate PSII quantum efficiency (Φ_II_) and linear electron flow (LEF) [34,35], as well as qL, a measure of the fraction of Q_A_ in the oxidised state [36], and the extent of NPQ based on the rapid “total” NPQ method developed by Tietz et al. [37], designated as NPQ_t_. Just prior to the saturation pulses, dark interval relaxation kinetics (DIRK, dark interval of approximately 300 ms) of the absorbance changes around 520 nm attributed to the electrochromic shift (ECS) were recorded. Fitting the ECS signals to exponential decay curves yielded estimates of the relative light-dark differences in thylakoid *pmf* (ECS_t_) and the proton conductivity of the chloroplast ATP synthase (*g*_H_+), as described in [16,38,39]. To account for differences in leaf thickness, light path or number of chloroplasts in various leaves, the ECS_t_ values were normalised to the relative chlorophyll contents as estimated by the SPAD parameter [33], which was measured at the end of the experiment. The extent of oxidation of P_700_ in the light was estimated by the DIRK of infrared LED light using an LED measuring pulse with peak emission at ∼810 nm.

### Environmental conditions during light potential measurements in the field

Supplemental figures S1A-C show the distributions of environmental factors (light intensities, leaf temperatures) for the measurements analyzed in this study. The MultispeQ sensor was positioned by the user to be parallel to the leaf surface, so that the cosine-corrected PAR sensor should effectively estimate PAR absorbed by the leaf surfaces *in situ* throughout their canopy, and thus the ambient PAR (PAR_amb_) values were dependent on both time of day (diurnal cycle, Fig. S1B) and by leaf angle (Fig. S1C). Ambient temperature and leaf temperatures (T_leaf_) were dependent on time of day, with obvious influences from weather-related fluctuations (Fig. S1A, B). We chose to compare results to T_leaf_, rather than ambient temperature, to better reflect the effects on leaf photosynthetic processes.

### Data calculations and cleaning

Data from the PhotosynQ platforms was reprocessed and cleaned to improve the estimation of decay constants for electrochromic shift and near infrared absorbance changes. As with any field experiments, some results were found to have obvious errors or be out of acceptable ranges, and were removed from the analysis. However, all original data was maintained in the online platform, allowing the reader to explore and reanalyse the effects of our data cleaning procedures. The rules and code for data flagging are defined in the Jupyter Notebook, (see Supplemental Information, “Data Cleaning Notebook”). A total of 292 points were flagged from a total of 1346 original measurements. The majority of the flagged measurements (179) were due to a defective device. The remaining 113 flagged points can be attributed to user error (e.g. leaf movements during measurements) or poor signal-to-noise that resulted in parameter values outside the theoretical ranges.

## Results

### Field measurements of photosynthetic parameters under ambient and rapidly elevated PAR

Figure 2A shows LEF measured at PAR_amb_ (LEF_amb_) plotted against ambient PAR_amb_ and leaf temperature (T_leaf_, see colouration of points). The plots use the square root of PAR to better resolve the results at lower PAR_amb_, and to partially linearise the responses. LEF_amb_ increased with increasing PAR_amb_, with a roughly hyperbolic dependence and an apparent half-saturation point of about 350 µmol photons m^-2^ s^-1^, reaching maximum values of about 250 µmol electrons m^-2^ s^-1^ at 1700 µmol photons m^-2^ s^-1^.

**Figure 2.**
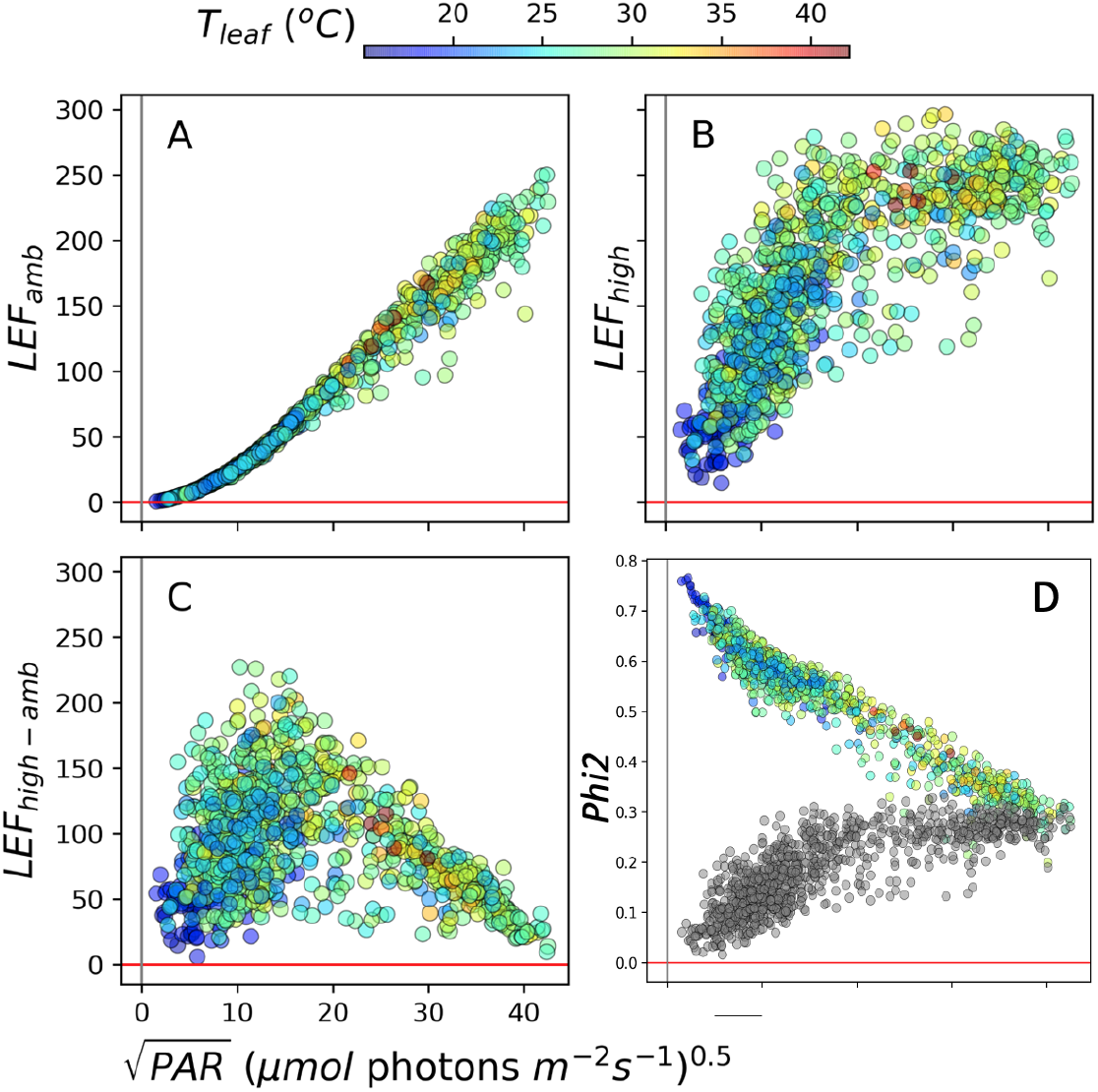
Light and temperature effects on linear electron flow (LEF) and Photosystem II quantum efficiency (Φ_II_). Each parameter was plotted functions of the square root of the ambient photosynthetically active radiation (PAR_amb_, X-axis) and Leaf Temperature (T_leaf_, coloration of points). A) Dependencies of Linear Electron Flow (LEF) measured at PAR_amb_; B) LEF measured at 10s high light (LEF_high_); C) The high light-induced differences in LEF (LEF_high-amb_); D) The PSII quantum efficiencies measured under ambient PAR (Phi2amb, points colored by T_leaf_) and at 10s high light (Phi2_high_, grey points).

Upon ten seconds of exposure to 2000 µmol photons m^-2^ s^-1^ increased LEF to generally higher values (LEF_high_ Fig. 2B), indicating that LEF_amb_ was at least partly light-limited under all of the conditions. Note that each LEF_amb_ point was taken on different leaves at different times (Materials and Methods) and has corresponding LEF_high_ and LEF_high-amb_ measurements. The relationship between measurements is illustrated in Fig. S2, which shows selected pairs of LEF_amb_ and LEF_high_ connected by vertical line segments. The extent of LEF_high_ was not uniform, but appeared to be strongly suppressed at low PAR_amb_ and/or low T_leaf_. The high light-induced difference in LEF (LEF_high-amb_) increased with PAR_amb_ at low light, reaching a peak at about 200 µmol photons m^-2^ s^-1^, above which it declined as PAR_amb_ approached PAR_high_ and LEF_high_ became light-saturated. The suppression of LEF_high_ was due to large decreases in the quantum efficiencies of PSII (Phi2, Fig. 2D). Phi2 at PAR_amb_ (Phi2_amb_) were highest at low PAR_amb_, and progressively saturated as light was increased. The opposite behavior was seen with Phi2 measured after 10s of high light (Phi2_high_ Fig. 2D, grey symbols) which was lowest at low PAR_amb_, and gradually increased with PAR_amb_.

### Gaussian Mixture Model clustering analysis of field data

A simple linear effects model applied over the entire data set (Supplemental Table S1A) indicated strong correlations between LEF_amb_ and both PAR_amb_ and T_leaf_, suggesting that both environmental factors controlled LEF_amb_. However, such correlations may be coincidental since PAR and T_leaf_ are both expected to be dependent on weather or time of day, as it is clear from the strong statistical correlations between PAR and T_leaf_. Also, the effects are likely to be co-dependent. For example, at low PAR_amb_, LEF_amb_ should be light-limited, and thus have minimal dependence on T_leaf_, but at higher PAR_amb_, may be more strongly controlled by temperature-dependent processes.

One approach to disentangling these effects would be to slice the data into segments, e.g., at different ranges of PAR_amb_, and test for correlations with T_leaf_ within each segment. However, arbitrary-chosen ranges for the segments can add bias, or fail to detect more complex interactions. We thus applied a Gaussian Mixture Model (GMM) clustering approach based on those presented earlier [40,41]. Because GMM is an unsupervised machine learning method, it can reduce bias in the selection of clusters that represent regions of distinct interactions among environmental and photosynthetic parameters. GMM assumes that the data points from the population of interest are being drawn from a combination (or mixture) of Gaussian distributions with certain parameters, and performs an optimization scheme to a sum of K Gaussian distributions, allowing for a completely unsupervised process, avoiding potential user bias. An expectation-maximization (EM) algorithm was used to fit the GMM to the dataset, generating a series of Gaussian components (clusters) with distributions characterised by specific means and covariance matrices. The optimal number of clusters was determined using the Bayesian Information Criterion (BIC), the value of the maximised log likelihood, with a penalty on the number of parameters in the model [40–43]. This approach also allows comparison of models with differing parameterizations and/or differing numbers of clusters, because the volumes, shapes, and orientations of the covariances can be constrained to those described by defined models [40].

Clusters obtained through GMM have both within cluster (intracluster) and between cluster (intercluster) variations. In order to test for intercluster variation, we used the clustering assignment obtained for one phenotype and applied it on other phenotype(s). Here we want to investigate what would be the distinctive behaviour of different phenotypes if we have used the same configuration. Using the same set of cluster assignments to different phenotypes, one might be skeptical of the clustering behaviour as phenotypes interact differently with PAR_amb_ and T_leaf_. In that case, we might not be able to directly compare the inter cluster behaviours of phenotypes. To mitigate this issue, we use the GMM clustering as a tool to create a “baseline” clustering configuration for one phenotype and use that configuration over another phenotype. We set up our hypothesis as two phenotypes are similar under the same configuration against they are not. If the interaction pattern of one phenotype with PAR_amb_ and T_leaf_ changes over the other phenotype, we reject our hypothesis and imply that different configurations of PAR_amb_ and T_leaf_ interact differently with phenotypes. By doing this we are able to disentangle the effect of PAR_amb_ and T_leaf_ and infer regarding the intracluster variations as to be a key element to determine variations in the interactions between parameters and variations in environmental conditions, e.g., to assess if a relationship is modulated in different ways under different ranges of conditions. Also, as will be seen in the Discussion, intercluster variations (differences in the mean and covariances between clusters) can be used to differentiate distinct patterns of behavior, or mechanistic interactions, between conditions.

As shown in Fig. S3, GMM analysis of LEF_amb_, PAR_amb_ and T_leaf,_ found six distinct, compact clusters that differed in the mode of interaction among the photosynthetic and environmental parameters. Encompassing points with lower PAR_amb_ showed strong (Clusters 1,2,4 and 5) dependence of LEF_amb_ on PAR_amb_, with little contributions from T_leaf_. By contrast, two clusters (3 and 6), which included points at higher PAR_amb_, showed substantial dependencies on both PAR_amb_ and T_leaf_. These results are consistent with LEF being predominantly light-limited at low ambient PAR, but increasingly limited by temperature-dependent processes at higher PAR. The presence of these two classes of clusters indicates that PAR_amb_ and T_leaf_ are likely to affect LEF_amb_ in independent ways. The fact that the shapes of the clusters were not determined with individual slicing under the individual parameters for PAR_amb_ and T_leaf_, but with a co-dependence on both PAR_amb_ and T_leaf_, suggests that, under some conditions, these effects interact, e.g. T_leaf_ may affect the dependence of LEF_amb_ on PAR_amb_.

GMM identified five distinct clusters for interactions among LEF_high_, PAR_amb_ and T_leaf_ (Fig. S4). In contrast to the results on LEF_amb_, clusters at lower PAR_amb_ (1, 2 and 4) showed LEF_high_ dependencies on both T_leaf_ and PAR_amb_, while Cluster 3 showed correlations with T_leaf_, but not with PAR_amb_. The stronger dependence on T_leaf_ of LEF_high_ compared to LEF_amb_ implies that the exposure to high light revealed additional rate limitations in LEF_high_ that were more strongly controlled by both T_leaf_ and PAR_amb_ and that, at least under some conditions, these effects were independent of each other.

### Analysis of NPQ

NPQ_t_ measured under PAR_amb_ (NPQ_amb_, Fig. 3A) showed a positive correlation to PAR_amb_, with an apparent tendency for smaller values at lower T_leaf_. NPQ_amb_ showed considerable variations, compared to LEF_amb_, even at low PAR_amb_, consistent with the idea that NPQ is governed not only by PAR but by metabolic, developmental or other environmental parameters.

**Figure 3.**
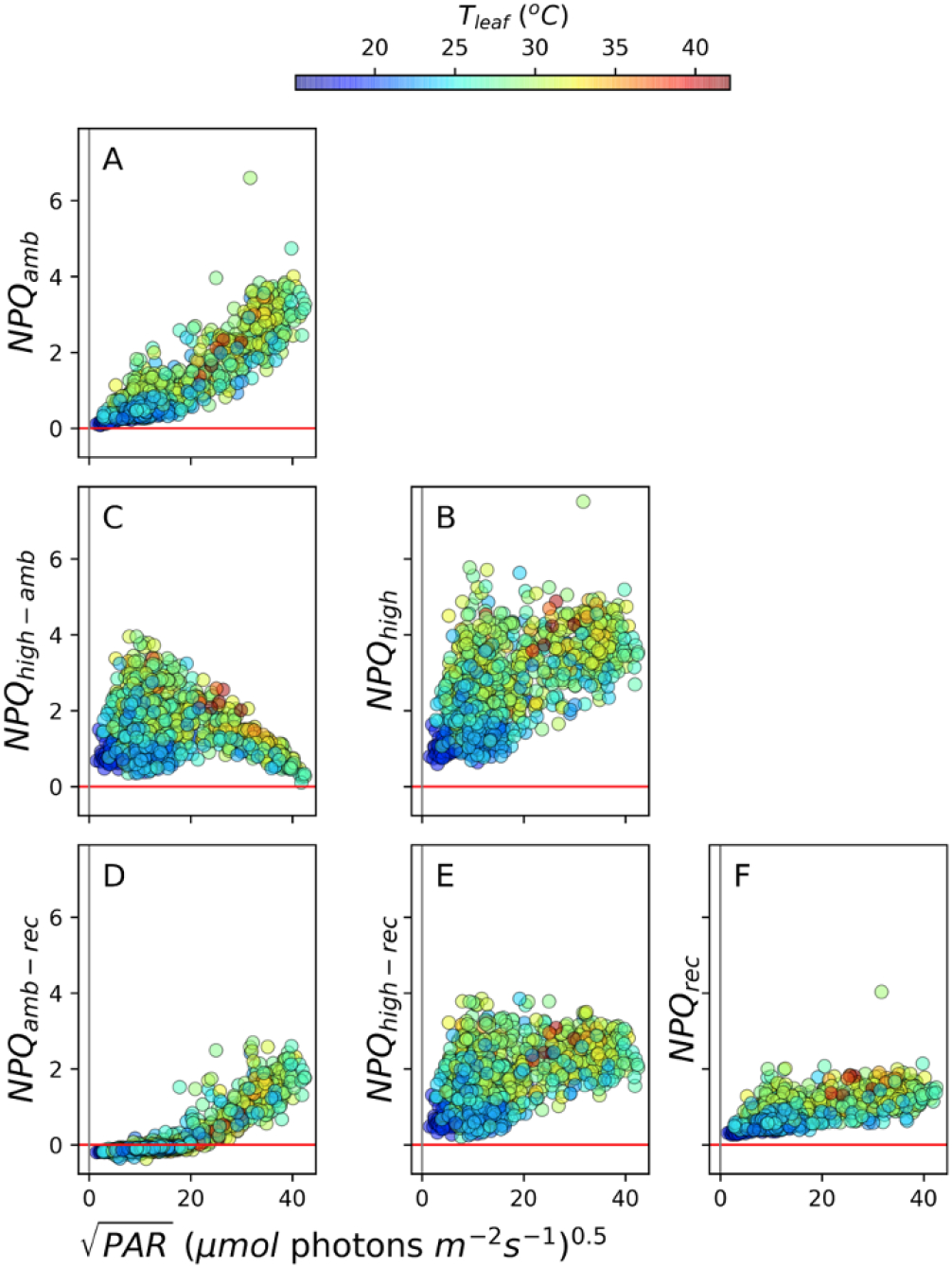
Light and temperature effects on NPQ. The NPQ parameter was plotted functions of the squar e root of the ambient photosynthetically active radiation (PAR_amb_, X-axis) and Leaf Temperature (T_leaf_, coloration of points). A) Induced NPQ measured at PAR_amb_; B) NPQ measured at 10s high light (NPQ_high_); C) The high light-induced differences in NPQ (NPQ_high-amb_); D) The difference in induced NPQ level at ambient PAR and the 10s recovery time in the dark (NPQamb_amb-rec_); E) The difference in induced NPQ level at 10s high PAR and the 10s recovery time in the dark (NPQ_high-rec_); F) the NPQ level after 10s in the dark (NPQ_rec_).

Figure 3B shows NPQ_t_ values measured at 10s full sunlight (NPQ_high_). The NPQ light potential, or light-induced differences in NPQ (NPQ_high-amb_) are shown in Fig. 3C. While NPQ_high-amb_ was always positive, both NPQ_high-amb_ and NPQ_high_ were suppressed at low PAR_amb_ or T_leaf_. NPQ_t_ measured after the 10s dark recovery period (NPQ_rec_, Fig. 3F) was consistently lower than NPQ_amb_ and NPQ_high_. The difference between NPQ_amb_ and NPQ_rec_ (NPQ_amb-rec_, Fig. 3D) ranged from slightly negative at low PAR_amb_, where the majority of NPQ_amb_ was rapidly reversible, to about one at the higher PAR_amb_, where about half of NPQ_amb_ was rapidly reversed.

Overall, these results indicate that the majority of NPQ_amb_ as well NPQ_high_ recovered within 10s of darkness and can likely be attributed to qE, and thus, under our conditions, qE is likely to be the most important form of NPQ. The residual, more slowly reversible, components reaching a little above 2 are likely to include qI or qZ [44,45], although the limited time frame for the protocol does not allow us to rule out contributions from longer-lived qE.

As with LEF, a simple linear effects model (Table S1B) showed strong interactions between T_leaf_ and PAR_amb_, on NPQ_amb_ and the corresponding GMM analysis identified four clusters (Fig. S5). Cluster 1, which encompassed the lowest range of PAR_amb_ values, showed strong dependence on PAR_amb_, with no significant dependence on T_leaf_. The remaining clusters showed either dependence solely on T_leaf_ (Cluster 4) or codependence on PAR_amb_ and T_leaf_ (Clusters 2 and 3). Because GMM clustering suggests that T_leaf_ and PAR_amb_ can interact or act independently, depending on conditions, we excluded the linear effects models and focused on GMM for analyses of the remaining parameters.

For the analysis of NPQ_high_ (Fig. S6), we used the clusters found for NPQ_amb_ (Fig. S5), allowing us to directly compare changes in correlations among parameters within each cluster [40]. Cluster 1, which encompassed the lowest range of PAR_amb_ values, showed strong dependence of NPQ_high_ on both PAR_amb_ and T_leaf_. This pattern of dependencies was in contrast to that for Cluster 1 for NPQ_amb_, which showed dependence solely on PAR_amb_. At a higher range of PAR_amb_ (Cluster 3), NPQ_high_ showed significant dependence solely on T_leaf_, again in contrast to the corresponding cluster for NPQ_amb_, which showed dependencies on both PAR_amb_ and T_leaf_. Overall, compared to NPQ_amb_, NPQ_high_ showed increased dependence on T_leaf_ in all clusters, suggesting that it is more substantially controlled by metabolic or physiological factors (see Discussion).

### The redox state of Q_A_

Figure 4A shows the dependencies of Q_A_ redox state (qL) on PAR and T_leaf_. qL measured at PAR_amb_ (qL_amb_, Fig. 4A), was relatively constant (ranging from about 0.3 to 0.75) across PAR_amb_, with somewhat higher values at both extremes of PAR_amb_. Lower leaf temperatures appeared to be associated with lower qL values, over the entire range of PAR_amb_, although the effect was particularly pronounced at low light. By contrast, qL measured at 10s of high light (qL_high_, Fig. 4B) showed strong dependence on PAR_amb_, ranging from near zero (fully reduced Q_A_) at low PAR_amb_, to almost one (fully oxidised) at higher PAR_amb_. Again, low T_leaf_ appeared to correlate with lower qL_high_ throughout the range of PAR_amb_. Strikingly, as shown in Fig. 4C, the high light treatment induced two distinct effects: At low PAR_amb_ and/or T_leaf_, it induced a net reduction of Q_A_, while it had the opposite effect at higher PAR_amb_ and T_leaf_. GMM clustering for qL_amb_, PAR_amb_ and T_leaf_ (Fig. S7) identified four distinct clusters. In Cluster 2, which encompasses points at low PAR_amb,_ significant associations were observed only between qL_amb_ and PAR_amb_. Clusters 1,3 and 4 (at higher PAR_amb_) showed co-dependencies between qL_amb_ and both PAR_amb_ and T_leaf_. GMM clustering for qL_high_, PAR_amb_ and T_leaf_ showed five distinct clusters (Fig. S8). Clusters 1,2 and 5, which encompassed generally lower ranges for PAR_amb_ and T_leaf_, showed qL_high_ dependencies on both PAR_amb_ and T_leaf_. Clusters 3 and 4 (generally with higher PAR_amb_ and T_leaf_ values) showed only dependencies on T_leaf_. The overall pattern of cluster behaviour was similar to that observed with respect to NPQ_amb_ and NPQ_high_.

**Figure 4.**
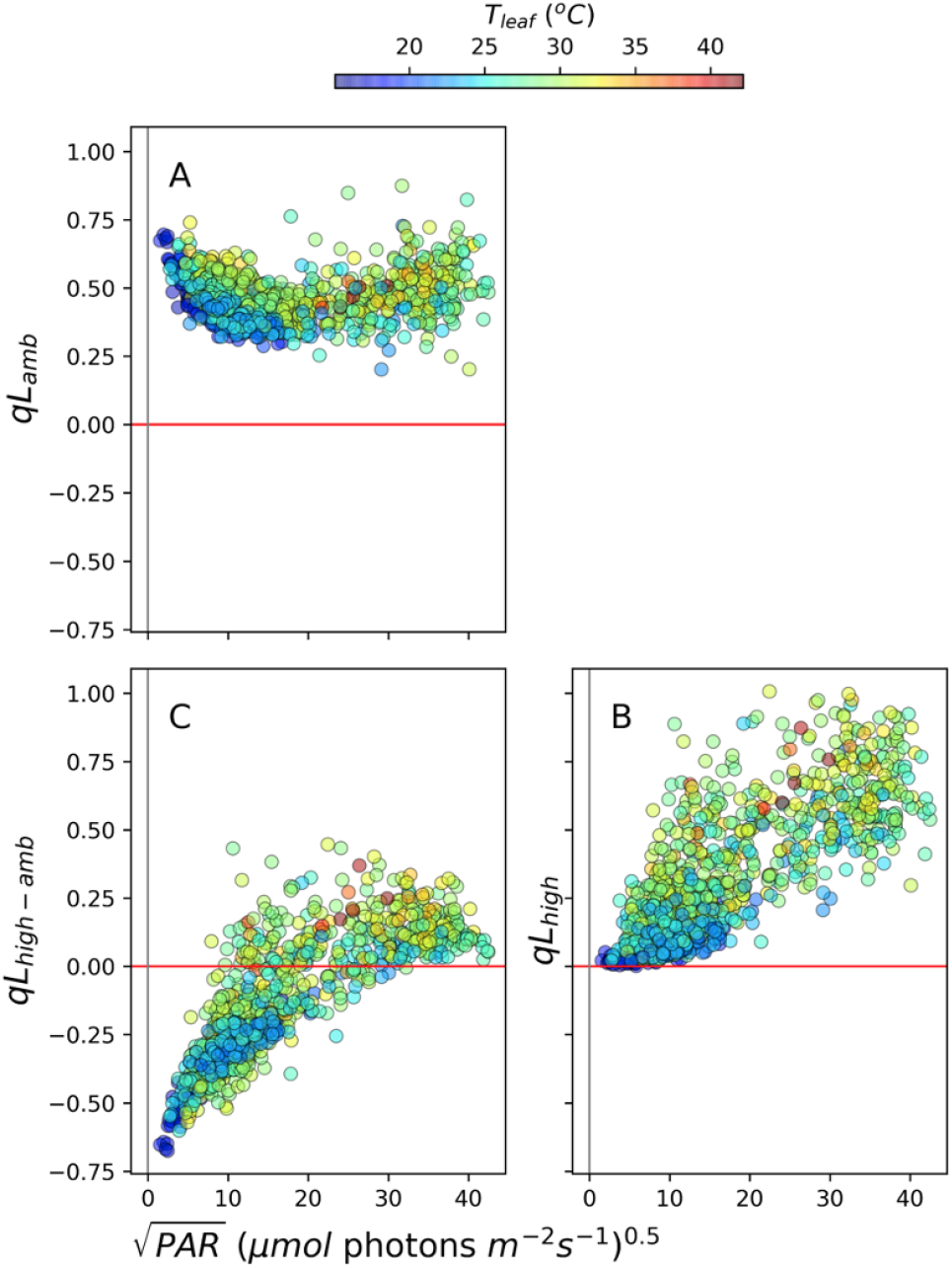
The light and temperature dependencies of the redox state of Q_A_. The qL parameter, a measure of fraction of Q_A_ in its oxidized state, was measured as described in Material and Methods, under ambient light (qL_amb_, Panel A), at 10s of high light (qL_high_, Panel B), and the change in qL between high and ambient PAR (qL_high-amb_, Panel C) as a functions of the square root of ambient PAR.

### P700 redox state

Figure 5 shows the extent of oxidised P_700_^+^ (P^+^), based on the DIRK of absorbance changes at 810 nm. P_700_^+^ at PAR_amb_ (P^+^_amb_, Fig. 5A), after ten seconds of high light (P^+^_high_, Fig. 5B) and the light-induced difference (P^+^_high-amb_, Fig. 5C). The extent of P^+^_amb_ was nearly linearly related to PAR_amb_. Increasing the light resulted in higher P^+^ values (P^+^_high_), indicating that, in all cases, PSI became more oxidised at high light. The extent of the light-induced oxidation was dependent on PAR_amb_, with lower extents at low PAR_amb_, and a peak at about 200-300 µmol photons m^-2^ s^-1^. The decrease at higher PAR_amb_ was probably due to the accumulation of pre-oxidised P_700_ prior to the high light treatment.

**Figure 5.**
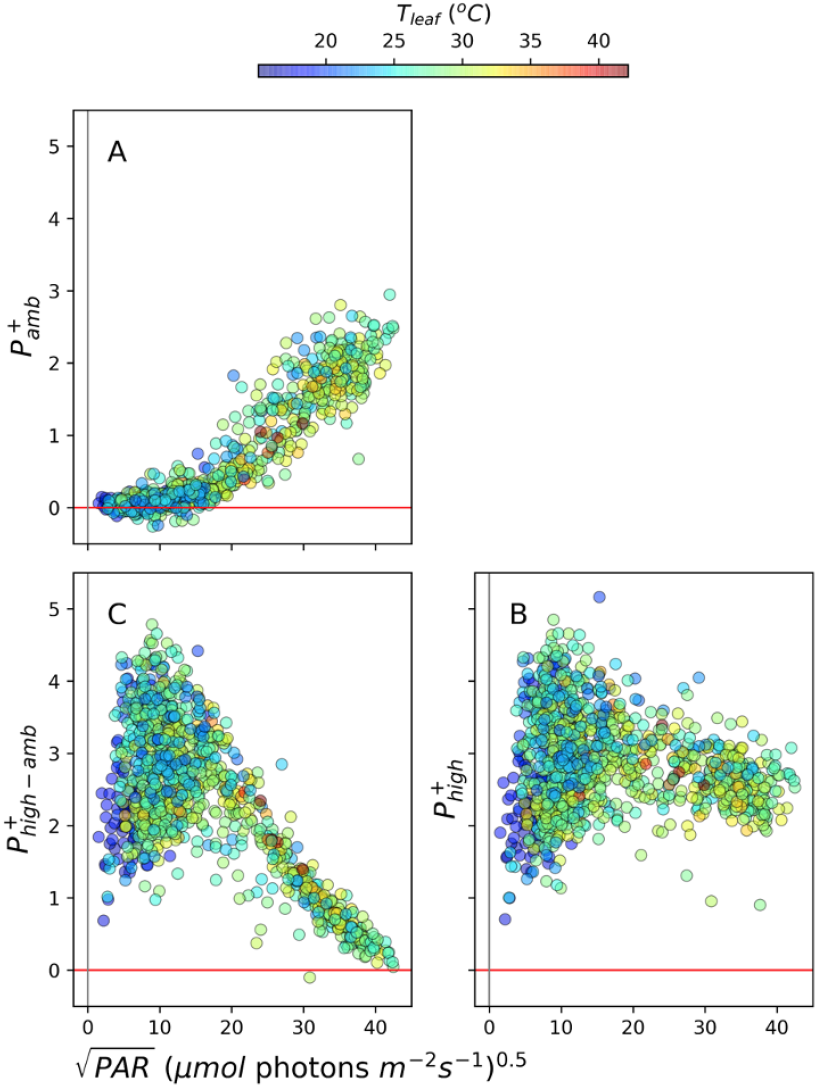
The light and temperature dependencies of the redox state of P_700_+. The redox state of P_700_ was measured using DIRK at 810 nm absorbance change under ambient light (P^+^_amb_, Panel A), at 10s of high light (P^+^_high_, Panel B), and the change in P^+^ between high and ambient PAR (P^+^_high-amb_, Panel C) as a functions of the square root of ambient PAR.

The full extent of P^+^_high_ was relatively constant over the conditions, suggesting that high light was able to nearly fully oxidise P_700_. However, there was a slight trend to lower P^+^_high_ at the highest PAR_amb_ or T_leaf_, suggesting that total oxidizable PSI may have decreased at high light or temperatures, perhaps reflecting accumulation of PSI photodamage or electron sink limitations. Consistent with these general trends, GMM analyses of P^+^_amb,_ PAR_amb_ and T_leaf_ identified four distinct clusters (Fig. S9), with dependencies on either PAR_amb_ by itself (Clusters 3 and 4), or both PAR_amb_ and T_leaf_ (Clusters 1 and 2). GMM clustering for P^+^_high_ identified five distinct clusters (Fig. S10), that showed a positive dependency of P^+^_high_ on either PAR_amb_ (Cluster 1), or T_leaf_ (Cluster 5), or a small, negative dependence on T_leaf_ (Cluster 3).

### ECSt and thylakoid *pmf*

Figure 6 shows dependencies of relative thylakoid *pmf*, estimated by normalised ECSt measurements, at ambient PAR (ECSt_amb_, Fig. 6A) and after 10s exposure to high light (ECSt_high_, Fig. 6B). The high light-induced differences (ECSt_high-amb_) are shown in Fig. 6C. ECSt_amb_ showed strong, positive correlations with PAR_amb_, similar to the responses of NPQ_amb_ (Fig. 3A) and P^+^_amb_ (Fig. 5A). ECSt_high_ values were, in general, larger than ECSt_amb_, resulting in positive values for ECSt_high-amb_. At low PAR_amb_, ECSt_high_ showed high variability, suggesting that the response is strongly dependent on other factors, but appeared to saturate (flatten) at higher PAR_amb_. These behaviours were reflected in ECSt_high-amb_, which showed strong variability at lower PAR_amb_ or T_leaf_, peaked at about 50-100 µmol photons m^-2^ s^-1^, and saturated at higher PAR_amb_.

**Figure 6.**
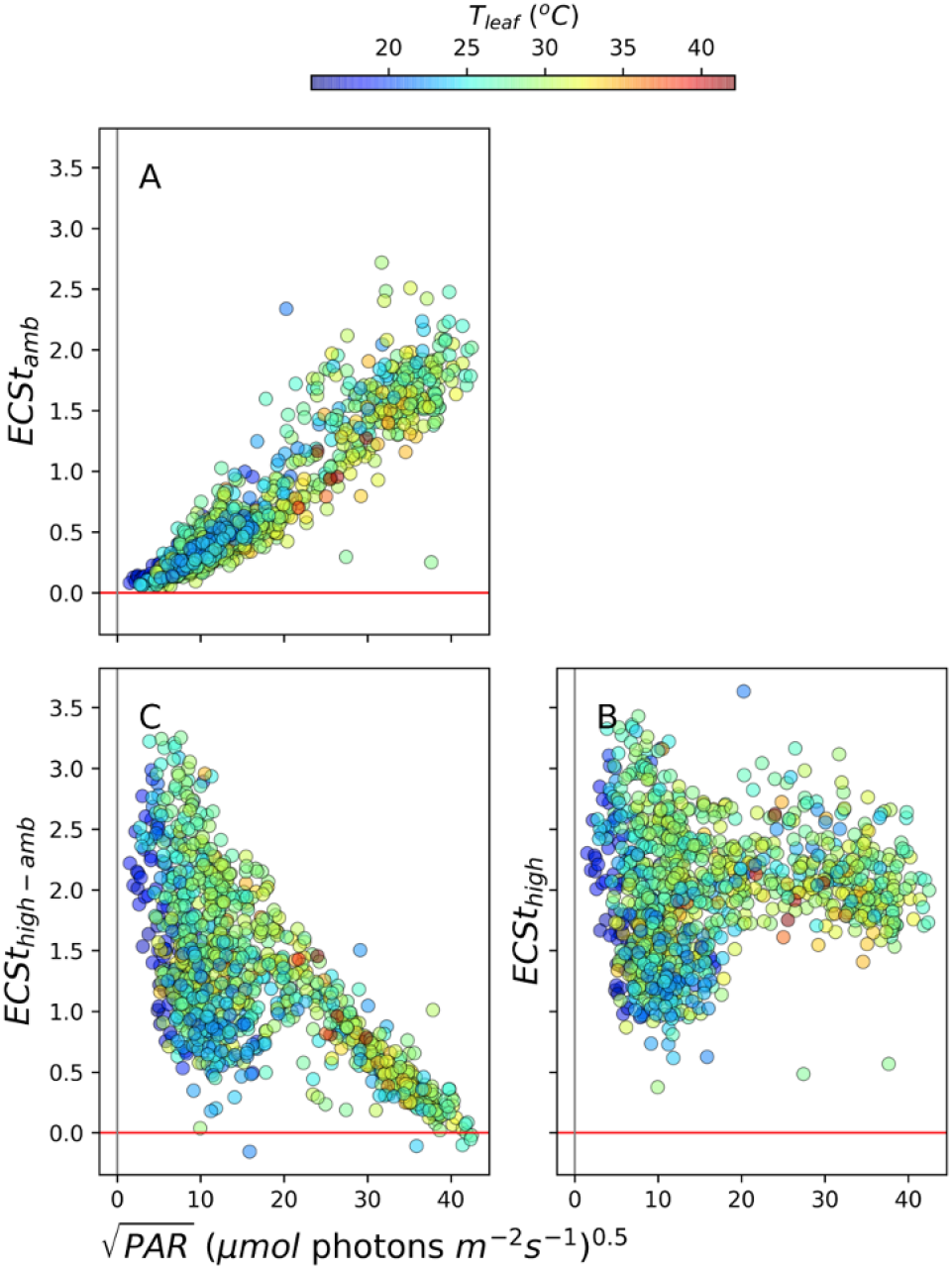
The light and temperature dependencies of the thylakoid *pmf* probed using ECSt signal. The *pmf* was measured using ECS under ambient light (ECSt_amb_, Panel A), at 10s of high light (ECSt_high_, Panel B), and the change in ECSt between high and ambient PAR (ECSt_high-amb_, Panel C) as a functions of the square root of ambient PAR.

GMM analysis of ECSt_amb_ identified five distinct clusters (Fig. S11). The cluster at the lowest range of PAR_amb_ (Cluster 1) showed dependence primarily on PAR_amb_. The remaining clusters showed positive correlations between ECSt_amb_ and PAR_amb_, but negative correlations with T_leaf_. By contrast, GMM of ECSt_high_ (Fig. S12) showed almost no dependence on either PAR_amb_ or T_leaf_, except at the lowest PAR_amb_ (Cluster 1) which showed negative correlations with PAR_amb_ and positive correlations with T_leaf_.

## Discussion

### Using PhotosynQ and MultispeQ to sample and resolve the effects of environmental fluctuations on photosynthetic processes

The MultispeQ measurements described above were designed to explore the photosynthetic responses of plants in a natural, fluctuating environment. In this type of field experiment, it is not possible to control all variables. Rather, the strategy was to “sample” responses under as many conditions as practical, while recording key metadata so that subsequent analyses can assess the impacts of various environmental fluctuations. Thus the observed trends may reflect both primary and acclimatory factors that change (or accumulate) over different time scales. Correlations that appear in such analyses can be used to test, at least to some extent, certain models, though it is important to note that more controlled experiments will be needed to fully determine cause-effect relationships, as discussed below.

A major outcome of the experiment is that, despite the fact that measurements were made over many plants, times etc., clear patterns of responses emerged that allow us to make some broad conclusions about the responses of photosynthesis to ambient and rapidly changing light. For example, the majority of NPQ_high_ was found, in general, to be rapidly variable, suggesting that qE was the major contributor: At lower PAR_amb_ that majority of NPQ_high_ was rapidly induced (see Fig. 3C), while at higher PARamb, pre-existing NPQ was rapidly recoverable (Fig. 3E) at higher PAR_amb_.

Another important trend was the suppression of the light potentials of both LEF (Fig. 2) and NPQ (Fig. 3) under some conditions, particularly under lower PAR_amb_ and/or T_leaf_. Further, strong decreases in LEF_high_ were not always accompanied by compensatory increases in NPQ_high_, implying that the productive and photoprotective light potentials can be simultaneously suppressed under certain conditions, a situation that is likely to promote the formation of reactive oxygen species and photodamage (see also below), with important implications for understanding the environmental robustness of photosynthesis [46].

### Disentangling interacting environmental impacts on photosynthetic processes

A key challenge to the field experiment approach is in teasing apart effects from different environmental factors, especially considering that such factors may be codependent or interact with each other in complex ways. For example, in visual inspection, most of parameters show apparent dependencies on both PAR_amb_ and T_leaf_ (e.g. Figs. 2-6) but, because increases in T_leaf_ are often correlated with increases in PAR_amb_, the effects of the two parameters may have been coincidental. It may also be that the environmental parameters interacted in complex ways, e.g. high PAR_amb_ may have exacerbated the effects of low T_leaf_. To address these issues, we applied an approach based on GMM to identify clusters representing distinct interactions among parameters. The approach is unsupervised, thus eliminating potential bias, while allowing us to test for changes in the environmental dependencies among multiple environmental parameters (Figs. S3-S12).

Analysis of GMM clusters implied that most parameters were dependent on both PAR_amb_ and T_leaf_, and at least under some conditions these effects are independent, or that one of the two factors predominates. Thus, the effects cannot be explained simply by coincidences between increased PAR and temperatures. Moreover, the non-rectilinear shapes of the clusters suggest that the effects of PAR_amb_ and T_leaf_ were interactive, e.g., changes in T_leaf_ modulated the effects of PAR_amb_ and vice versa. Overall, these interactions are in line with well-known temperature and PAR dependence of photosynthesis, but this type of analyses can reveal the specific combination of conditions that induce distinct behaviours, allowing for assessments of the involvement of specific mechanisms (see below) and to identify genotypic or management impacts on crop resilience and productivity.

At low PAR_amb_, we expect steady-state photosynthesis to be predominantly light-limited, and thus the effects of T_leaf_ should be low. As light increases, downstream biochemistry should become increasingly limiting. Because downstream energy storage and metabolic processes are likely to be more temperature dependent than photochemistry, this shift may allow us to distinguish between these types of limitations. Such behaviours are apparent in many of the measured parameters, e.g., LEF_amb_, which was not substantially dependent on T_leaf_ at low PAR_amb_, but became codependent on PAR_amb_ and T_leaf_ at higher PAR_amb_ (Figs. 2A, S3), consistent with a progressive shift from light-limitation to assimilation-limitation. Similarly, NPQ_amb_ was solely dependent on PAR_amb_ in the cluster at low PAR_amb_, but became increasingly dependent on T_leaf_ as PAR_amb_ increased (Fig. 3A). This shift is consistent with a control of NPQ_amb_ by PAR (at low PAR_amb_) and downstream metabolic processes, particularly at higher PAR_amb_, e.g., due to regulation of the ATP synthase activity or cyclic electron flow (CEF) [47].

By contrast, LEF_high_ and NPQ_high_ showed much greater dependence on T_leaf_, and these differences were more pronounced when the high light was imposed on leaves at low PAR_amb_ and T_leaf_, i.e. the opposite of what was seen for LEF_amb_ and NPQ_amb_. Interestingly, the LEF_high_ rates achieved in leaves exposed to lower PAR_amb_ were strongly suppressed below the maximum LEF_amb_ values measured at higher PAR_amb_ (compare Figs. 2A and B), This behavior suggests that the suppression of LEF_high_ occurs when abrupt increases in light overwhelm the activation of downstream energy storage and metabolic processes. This is generally consistent with observations that the activities of metabolic enzymes are regulated to match the availability of energy from the light reactions, which involve large suite of co-regulatory processes, as extensively reviewed elsewhere, e.g. [16,47–53], but that these responses lag behind the changes in light. The *in situ* light potential measurements afforded by MultispeQ show that these situations are very likely to occur under many field situations.

These results also imply that accurate estimates of LEF, NPQ and other photosynthetic parameters will require measurements under ambient light, because sudden changes in PAR can lead to severe perturbations in photosynthetic limitations or regulation. Attempts to “simplify’’ field experiments by setting PAR to some constant value will lead to strong artifacts. Such effects are vividly demonstrated by the opposite dependencies of Phi2_amb_ and Phi2_high_ on PAR_amb_ (Fig. 2D), and validate the use of the PAR matching feature of the MultispeQ instrument. It is important to keep in mind that the rates of acclimation may vary substantially between species, and that these may be assessed by performing more intensive experiments with variable high light and dark recovery times.

### Mechanisms for controlling the light potentials of LEF and NPQ using MultispeQ field data

The rapid reversal of NPQ_amb_ and NPQ_high_ over 10 s of dark indicated that, under our conditions, NPQ is predominantly in the form of qE (Fig. 3B and C), and thus dependent on lumen acidification and subsequent pH-dependent responses. Lumen acidification can be controlled by changes in proton influx (through changes in LEF and CEF), proton efflux through the ATP synthase and the partitioning of *pmf* into electric field (*Δ*ψ) and *Δ*pH components, which in turn, are impacted by metabolic status, as proposed earlier [15,38]. Here, we explore the possible mechanistic bases for these effects, by comparing the correlations among MultispeQ measurements.

Scheme 1 illustrates three basic mechanistic models describing proposed processes that can limit the light potentials of photosynthetic and photoprotective mechanisms. The models make qualitative predictions about how the actions of each mechanistic model will impact correlations between measured photosynthetic parameters, and thus can be used as a framework for interpreting the field data introduced in Results. The expected effects on the measured parameters are summarised in Scheme 1, which shows specific effects of each model.

**Scheme 1:**
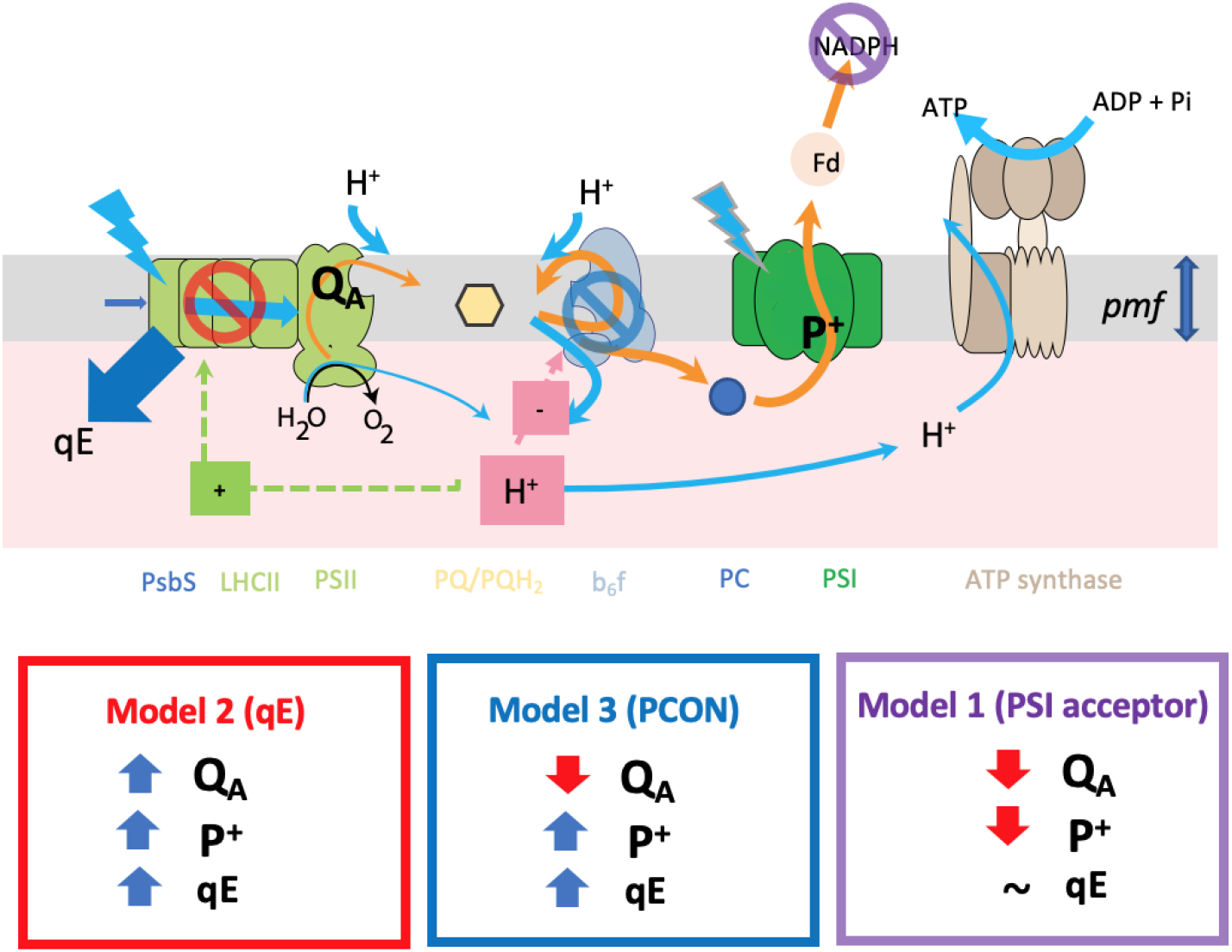
Models for limitations to LEF and NPQ light potentials.

**Model 1: PSI acceptor-side limitations** (Scheme 1, Model 1) where lack of NADP^+^, ferredoxin or other PSI acceptors prevent further LEF. We expect this limitation to result in accumulation of electrons throughout the electron transfer chain, thus resulting in net reduction of Q_A_ (decreasing qL) and P_700_^+^ (decreasing the 810 nm absorbance signal). The decreases in proton fluxes associated with backup of electrons may, in addition, prohibit rapid, light-induced increases in *pmf*, lumen acidification and qE responses.

**Model 2: Increased NPQ** (Scheme 1, Model 2). Increased NPQ should decrease delivery of excitation energy to PSII (but not to PSI), resulting in net oxidation of Q_A_ (increasing qL) and P_700_^+^ (increased 810 nm DIRK signal). Under some conditions, the NPQ will be rapidly induced by increased *pmf* and lumen acidification followed by activation of qE, which should be visible as increased NPQ_high-amb_. Under other conditions, e.g. at higher PAR_amb_, NPQ may already have been induced. If this NPQ is in the form of rapidly-reversible qE, it should substantially decay during the 10-s dark recovery period, resulting in increased NPQ_high-rec_. More slowly-induced or relaxing forms of NPQ, including qI, qZ and long-lived qE, may be also present prior to and throughout the experiment. The forms should register as increases in NPQ_rec_, but not in NPQ_high-amb_ or NPQ_high-rec_, but given that the high light and recovery periods were only 10s long, our results do not allow us to distinguish among these possible forms.

**Model 3: Photosynthetic control (PCON**, Scheme 1, Model 3). PCON results from the slowing of PQH_2_ oxidation at the cytochrome *b*_*6*_*f* complex as the lumen becomes acidified. If PCON occurs without activation of qE, we expect a net reduction of Q_A_ (decreasing qL) but a net oxidation of P^+^ (increasing the 810 nm absorbance signal).

The qE and PCON models can be further subdivided [18,54]. In most cases, we expect lumen acidification accompanied by elevated *pmf*, reflected in an increased ECSt signal, which can be induced by increased proton influx into the lumen, due to increased LEF, increased CEF, or decreased conductivity of the thylakoid to protons (*g*_H_^+^) by slowing the ATP synthase. Alternatively, lumen acidification can also be associated with an increase in the fraction of *pmf* that is stored as ΔpH, by controlling the flow of counterions across the thylakoid membrane, altering the partitioning of *pmf* in ΔpH and Δ*ψ*. In this case, acidification may occur with little or no increases in total *pmf*, or the rates of proton influx, though the current field-based data do not allow us to directly distinguish these possibilities.

These models, while not mutually exclusive, will tend to counteract each other, at least within a particular leaf. For instance, PSI acceptor side limitations will tend to inhibit electron flow, thus decreasing proton flux and *pmf* generations. On the other hand, the generation of *pmf* will tend to slow electron flow (through Models 2 or 3), thus preventing the buildup of electrons on PSI electron acceptors. However, it is important to note that, in a survey type experiment like ours, photosynthesis in different leaves may be limited by distinct processes, and thus any collection of samples may reflect various combinations of the above models.

### Testing models for limitations in light potentials

By plotting MultispeQ parameters against each other, we can test for more detailed patterns of behaviours predicted by the above models. Figure 7 shows that P^+^_high-amb_ (high-light-induced P_700_ oxidation) was positively correlated with light-induced increases in *pmf* (ECSt_high-amb_). Under all conditions, increasing PAR from PAR_amb_ to PAR_high_ resulted in a net oxidation of P_700_, i.e., P^+^_high-amb_ was consistently positive. This behaviour is consistent with Models 2 (NPQ) or 3 (PCON), both of which predict a decrease in delivery of electrons from PSII to PSI. By contrast, we did not see evidence for high light-induced net *reduction* of P_700_^+^, i.e., values of negative P^+^_high-amb_, implying that Model 1 was not a major limitation to light potential. This does not exclude Model 1 from limiting photosynthesis in different species and conditions, as has been proposed to be important in chilling sensitive plants [55]. However, the apparent avoidance of Model 1 (or prevalence of Models 2 and 3) behaviour may reflect the “tuning” of the light reactions to prevent the accumulation of reduced electron acceptors of PSI associated with photodamage [23], and the associated O_2_ caused by buildup of electrons on PSI [56].

**Figure 7.**
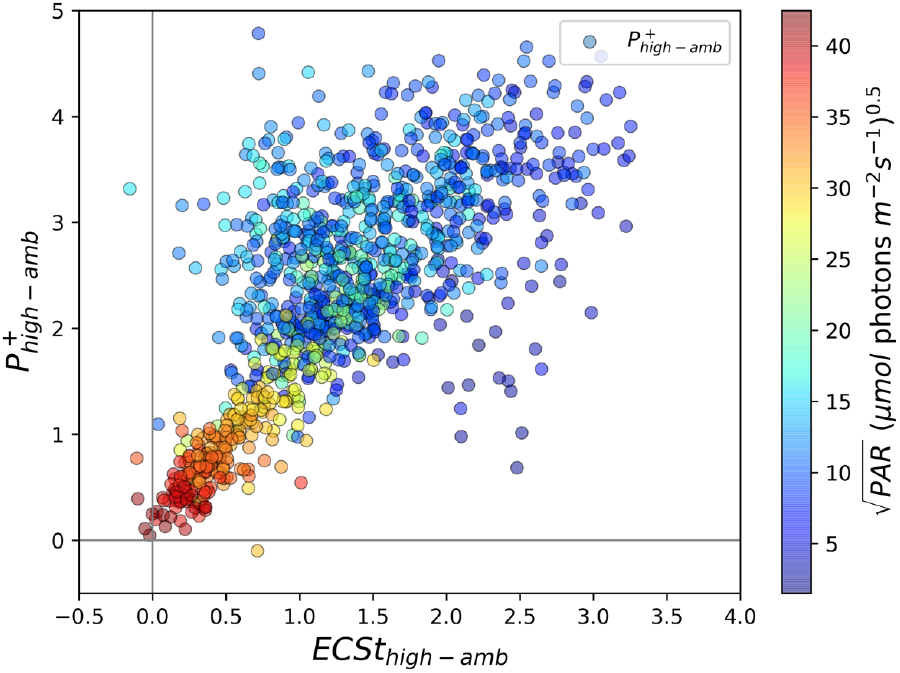
The relationship between light-induced thylakoid *pmf* and changes in P_700_ redox state. Changes in the thylakoid pmf (ECSt_high-amb_) were estimated using the ECSt parameter, and changes in P_700_^+^ were measured using the absorbance changes at 810 nm, as described in Materials and Methods, under ambient light (ECSt_amb_, P^+^_amb_) and after 10s of high light (ECSt_high_ P^+^_amb_). The coloration of the points was set to a function of the square root of ambient PAR (PAR_amb_).

Overall, the behaviours seen in Fig. 7 are consistent with restrictions in electron flow to PSI imposed by increases in *pmf*, most likely through the acidification of the thylakoid lumen. In the case of Model 2 (rapid NPQ), this would be related to the induction of qE, while in Model 3 (PCON), this could be related to slowing of electron flow at the cytochrome *b*_*6*_*f* complex.

Figure 8A further investigates this behaviour by plotting the dependence of high light-induced changes in P_700_^+^ (P^+^_high-amb_) with changes in Q_A_ redox state (qL_high-amb_). The expected theoretical behaviours for the three models are indicated by the coloured boxes in the figure, and can be related to Models 1-3 in Scheme 1:

**Figure 8.**
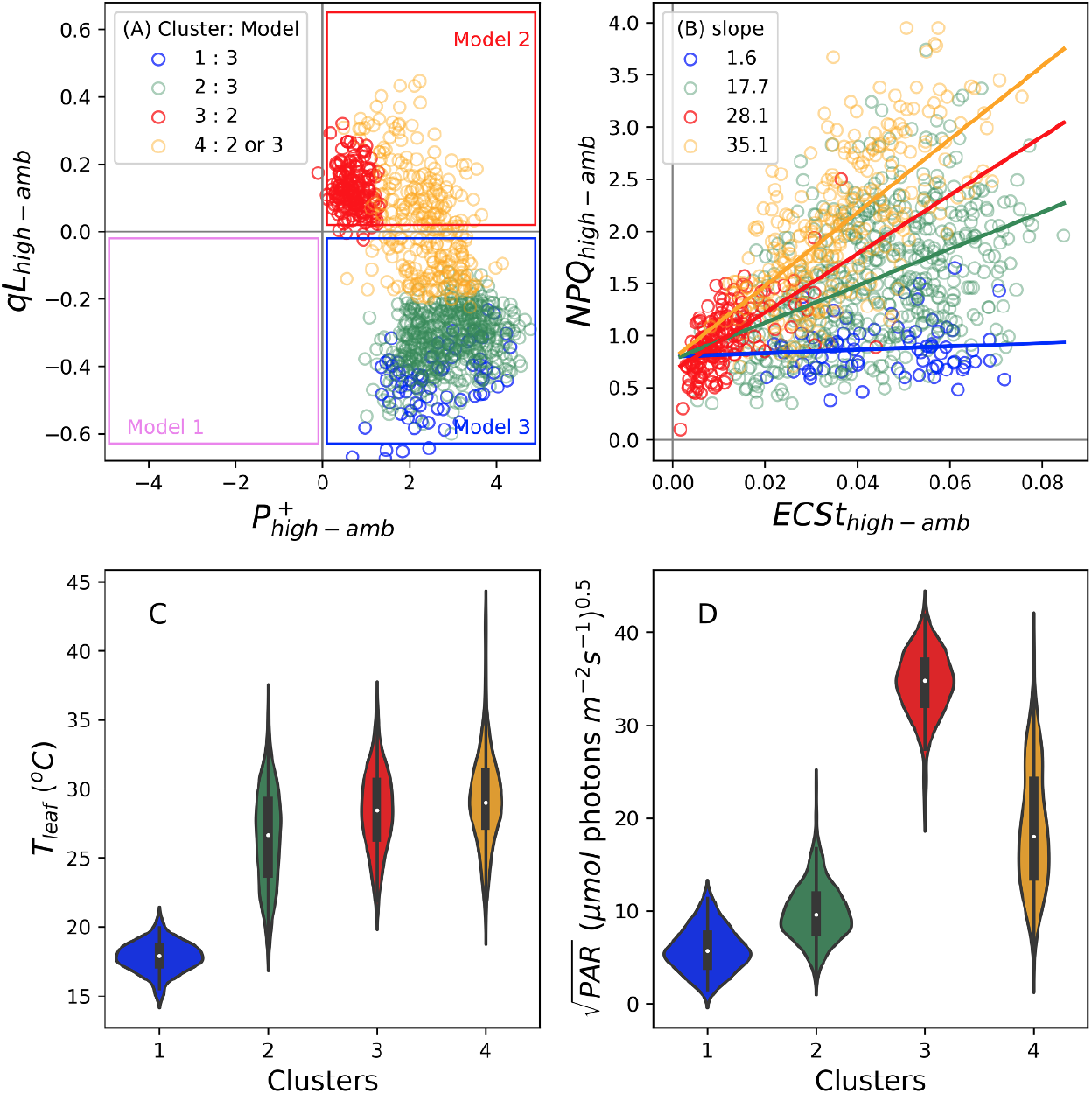
The relationships between light-induced changes in Q_A_ redox state and P_700_ redox state (Panel A), and between rapidly inducible NPQ and thylakoid *pmf* (Panel B) and the leaf temperature (Panel C) and PAR (Panel D) dependencies of Gaussian Mixture Models (GMM) clusters. Changes in P_700_^+^ (P^+^_high-amb_), Q_A_ redox state (qL _high-amb_), rapid changes in NPQ (NPQ _high-amb_) and thylakpoid *pmf* (ECSt_high-amb_) were measured as described in Materials and Methods. Data were clustered using the GMM approach described in the text, resulting in four distinct clusters, designated buy the blue, green, red and ocher symbol colors (see legend in Panel A). In Panel B, the slopes for the relationship between NPQ_high-amb_ and ECSt_high-amb_ were estimated by linear regression (slopes for clusters 1,2,3 and 4 were estimated to be 1.6, 17.7,28.1 and 35.1, respectively). Panels C and D show distributions of leaf temperatures (T_leaf_, Panel C) and square root of ambient PAR for each cluster in Panels A and B.

- **Model 1** (violet box) predicts net **reduction** of P_700_ (P^+^_high-amb_ < 0) and net **reduction** Q_A_ (qL_high-amb_ < 0)
- **Model 2** (red box) predicts net **oxidation** of P_700_ (P^+^_high-amb_ >0) and net **oxidation** Q_A_ (qL_high-amb_ > 0)
- **Model 3** (blue box) predicts net **oxidation** of P_700_ (P^+^_high-amb_ > 0) but net **reduction** Q_A_ (qL_high-amb_ < 0)

We observe behaviours consistent with both Models 2 and 3, suggesting that the behaviour of the system changed with conditions. Note that the boxes in Figure 8A represent “pure” behaviours, and it is possible that the effects of a particular mechanism may be intermediate, e.g., the responses may be limited by a combination of reduction of Q_A_ and increased NPQ.

Figure 8B plots the dependence of NPQ_high-amb_, which can be attributed to light-induced qE changes, on light-induced *pmf* changes (ECSt_high-amb_). A generally positive correlation was observed between NPQ_high-amb_ and ECSt_high-amb_, but with high variability, especially at higher values. Applying the clustering obtained for Fig. 8A on top of the data in Fig. 8B, we see that this variability can be explained by the environmental conditions and the modes of behaviours. Specifically, we see clear evidence for condition-dependent suppression of rapid activation of qE in response to increases in *pmf*. Particularly, the sensitivities of NPQ_high-amb_ to ECSt_high-amb_, as indicated by the slopes in Fig. 8B, were smallest in Clusters 1 (slope ∼ 1.6) and 2 (slope ∼ 17.7), which comprise those with Model 3-like behaviour and occured at low T_leaf_ and PAR_amb_ values. Higher sensitivities of NPQ_high-amb_ to ECSt_high-amb_ were seen for Clusters 3 (slope ∼ 28.1) and 4 (slope ∼35.1), which comprised those associated with Models 2 and intermediate, and occurred at higher T_leaf_ and PAR_amb_ values.

To assess what controlled the switch between Models 2 and 3, we performed GMM (using qL_high-amb_, P^+^_high-amb_, T_leaf_ as inputs). Four distinct clusters were observed (see symbol colours, Fig. 8A). Intercluster comparisons show that points in Clusters 1 and 2 fell exclusively in the region predicted for Model 3. Cluster 3 fell entirely within the region predicted for Model 2. Cluster 4 extended between these regions, possibly indicating contributions from both mechanisms. The clusters falling in the Model 3 region were associated with relatively low T_leaf_ (Fig. 8C) and PAR_amb_ (Fig. 8D), compared to those associated with Model 2 or intermediate behaviours, suggesting that Model 2 prevailed at higher T_leaf_ and/or PAR_amb_, while Model 3 prevailed at lower values. Within the GMM clusters (Fig. S13), qL_high-amb_ was dependent predominantly on T_leaf_ (Cluster 3), PAR_amb_ (Cluster 1), or both (Clusters 2 and 4). This dependence suggests that T_leaf_ and PAR_amb_ acted either independently or cooperatively, depending on conditions, affecting the propensity for photosynthesis to adopt Model 2 or 3 behaviours.

The data in Figs. 8 show that, at lower T_leaf_ and PAR_amb_, qE activation was suppressed despite light-induced increases in *pmf*, and that this behaviour was associated with accumulation of electrons on Q_A_ but oxidation of P_700_ (Fig. 8A), suggesting that, under these conditions, light-induced increases in *Δ*pH caused slowing of the cytochrome *b*_*6*_*f* complex (PCON), but that the qE response lagged behind or was completely suppressed, leading to Model 3 behaviour.

It has been shown that the lumen pH-dependencies of qE and PQH_2_ oxidation by the cytochrome *b*_*6*_*f* complex are tightly coordinated, so that increased lumen acidity activates photoprotection prior to PCON, presumably to prevent the accumulation of reduced Q_A_ [54]. However, these experiments were performed under more slowly-changing (near steady-state) conditions in the laboratory, and our results suggest that this coordination can be defeated under real world conditions in the field, especially when T_leaf_ is low and PAR fluctuates rapidly. This discoordination can have strong implications for photodamage, as it has been shown that high thylakoid *pmf* can greatly accelerate PSII recombination reactions, especially when Q_A_ is reduced, leading to ^1^O_2_ production [28,29]. It thus seems reasonable to suggest that the shift from qE to PCON at low T_leaf_ will increase the rates of photodamage.

There are several possible mechanisms by which the response of qE can be uncoupled from increased *pmf*. Longer-term dependencies of NPQ on temperature have been reported under both field [57–59] and laboratory [60][61] conditions. The current work shows effects on rapid NPQ and LEF changes, which can be related to distinct mechanistic models. For example, it is known that the xanthophyll cycle is strongly temperature dependent, though the general observation is that zeaxanthin tends to accumulate at lower temperatures due to a slowing of the epoxidation of zeaxanthin [60][61,62]. Interestingly, we would expect the accumulation of zeaxanthin to augment, rather than suppress qE responses as we have observed in the current results. Lumen acidification may also be rate limiting for formation of qE. While rapid increase in light can result in nearly instantaneous increases in *Δ*ψ, formation of *Δ*pH and lumen acidification requires counterion transport processes, which tend to be slow, and thus lumen acidification lags behind [25,29], and it is possible that this process is substantially slowed at low temperature. Other possible limitations include temperature-dependence of conformational rearrangement of antenna complexes following protonation of PsbS [63,64], which in turn may be related to the interactions among thylakoid proteins, lipids and ultrastructure [12,44,65,66]. The current data does not allow us to discriminate between these models with the current data set, but the work suggests conditions and species under which such limitations occur, and how they may impact plant productivity or resilience.

### Conclusion: Current limitations and prospects for open science-led efforts to understand and improve photosynthesis

There are intense, ongoing efforts to improve photosynthesis, yet the importance of the responses of photosynthesis under fluctuating, real world conditions are just now being recognised. In particular, we lack understanding of the extents and impacts of these responses, as well as their mechanisms and genomic control, which will be critical to achieving field-relevant improvements in efficiency and robustness, especially in a changing environment.

Here, we demonstrate methods and tools to assess the light responses of photosynthetic processes under real world conditions, and use them to explore the factors that limit the capacity of plants to utilise or dissipate rapidly increased PAR. A major outcome is that, despite the complexities of field environments, clear behavioural patterns can be resolved, as long as the experiment contains a sufficient number of points taken over a large environmental space, and that includes both environmental metadata. Such combinations of information allow for the generation and testing of specific hypotheses. For example, we observed no evidence for Model 1 behaviour in the current study, but we do not exclude the possibility in different species and/or different environmental conditions. The rapid measurements allowed us to test for various models over broad-scales by looking for internally-consistent relationships among the various measured parameters. Further, while we surmised (above) that Model 3 type behaviour would likely lead to photodamage, we do not have independent endpoint measurements (e.g., yield, growth rates etc.) to validate that the propensity for Model 3 behaviour has long-term consequences. The models are not exclusive, and there will almost certainly be cases, e.g., Cluster 4 in Fig. 8, where intermediate behaviours will be apparent, either because of co-limitations among multiple processes or heterogeneity between chloroplasts in the leaf samples.

We also emphasise that the data presented here was intended to introduce the approaches and methods, and thus leaves a number of questions unanswered, but sets up the approach to further study. The origins of these effects may include several classes of processes [31,67], that may differ under different conditions [68], including induction of downstream assimilatory reactions and metabolic pools [69,70], downstream sink reactions [71], redox regulation [72,73], balancing between the production and consumption of ATP and NADPH [1,49], ion homeostasis and regulation of thylakoid *pmf* [25,74], low stomatal aperture that may lead to transient depletion of internal CO_2_ levels. Distinguishing these will likely require more detailed phenotyping and biochemical [10,56,75], modeling [31] and genomics and genetics approaches [76].

The accessibility of the tools should allow larger numbers of researchers to answer these types of questions over a broader set of results, as will be presented in an upcoming paper. This approach was made possible by the combination of several open science advances. Collation of large amounts of data and metadata through the MultispeQ and PhotosynQ platforms [33], allowing us to explore the interdependencies of multiple phenotypes and environmental conditions (metadata). The GMM methods allowed us to explore the interactions among multiple environmental parameters and photosynthetic phenotypes, and test for the participation of distinct mechanistic models to explain the limitations to photosynthesis under field conditions, leading to the identification of distinct limitations in the rapid activation of NPQ and LEF at low temperature. Finally, making all tools, protocols and analytical methods available in directly usable forms, the project can be readily expanded to include multiple environments and species, as well as alternative models.

## Acknowledgements

The authors thank Dr. Ute Armbruster, Thekla von Bismarck, Dr. Nicholas Fisher, Dr. Jennifer Johnson, Dr. Thomas Avenson and Oliver Tessmer for valuable discussions and critical reading of the manuscript and for numerous contributors to PhotosynQ data sets.

## Data accessibility

**Primary data is available on the** photosynq.org site under the project “rapid-ps-responses-pam-ecst-npqt-mint-dmk”. Data cleaning and analysis code is available in a GitHub repository (https://github.com/protonzilla/Light-Potentials-in-Field).

## Author contributions

A.K. and D.M.K. designed the experiments. A.K. and H.T. conducted experiments. A.K., A.C., S.K. and D.M.K. analyzed data. A.K., A.C., S.K., T.M. and D.M.K. contributed to the interpretations of data and writing manuscript.

## References

1. Kramer DM, Evans JR. 2011 The importance of energy balance in improving photosynthetic productivity. Plant Physiol. 155, 70–78.

2. Li XP, Gilmore AM, Caffarri S, Bassi R, Golan T, Kramer D, Niyogi KK. 2004 Regulation of photosynthetic light harvesting involves intrathylakoid lumen pH sensing by the PsbS protein. J. Biol. Chem. 279, 22866–22874.

3. Kromdijk J, Glowacka K, Leonelli L, Gabilly ST, Iwai M, Niyogi KK, Long SP. 2016 Improving photosynthesis and crop productivity by accelerating recovery from photoprotection. Science 354, 857–861.

4. Li XP, Muller-Moule P, Gilmore AM, Niyogi KK. 2002 PsbS-dependent enhancement of feedback de-excitation protects photosystem II from photoinhibition. Proc. Natl. Acad. Sci. U. S. A. 99, 15222–15227.

5. Eskling M, Arvidsson P-O, Akerlund H-E. 1997 The xanthophyll cycle, its regulation and components. Physiol. Plant. 100, 806–816.

6. Demmig-Adams B. 1990 Carotenoids and photoprotection in plants: A role for the xanthophyll zeaxanthin. Biochim. Biophys. Acta 1020, 1–24.

7. Niyogi KK, Björkman O, Grossman AR. 1997 The roles of specific xanthophylls in photoprotection. Proc. Natl. Acad. Sci. U. S. A. 94, 14162–14167.

8. Eskling M, Emanuelsson A, Akerlund H-E. 2001 Enzymes and mechanisms for violaxanthin-zeaxanthin conversion. In Regulation of Photosynthesis (eds E-M Aro, B Anderson), pp. 806–816. Dordrecht, The Netherlands: Kluwer Academic Publishers.

9. Kunz HH, Gierth M, Herdean A, Satoh-Cruz M, Kramer DM, Spetea C, Schroeder JI. 2014 Plastidial transporters KEA1, -2, and -3 are essential for chloroplast osmoregulation, integrity, and pH regulation in Arabidopsis. Proc. Natl. Acad. Sci. U. S. A. 111, 7480–7485.

10. Armbruster U et al. 2014 Ion antiport accelerates photosynthetic acclimation in fluctuating light environments. Nat. Commun. 5, 5439.

11. Davis GA, William Rutherford A, Kramer DM. 2017 Hacking the thylakoid proton motive force (pmf) for improved photosynthesis: Possibilities and pitfalls. Philos. Trans. R. Soc. Lond. B Biol. Sci. 372, 20160381.

12. Jahns P, Holzwarth AR. 2012 The role of the xanthophyll cycle and of lutein in photoprotection of photosystem II. Biochim. Biophys. Acta 1817, 182–193.

13. Malnoë A, Schultink A, Shahrasbi S, Rumeau D, Havaux M, Niyogi KK. 2018 The Plastid Lipocalin LCNP Is Required for Sustained Photoprotective Energy Dissipation in Arabidopsis. Plant Cell 30, 196–208.

14. Schiphorst C, Bassi R. 2020 Chlorophyll-Xanthophyll Antenna Complexes: In Between Light Harvesting and Energy Dissipation. In Photosynthesis in Algae: Biochemical and Physiological Mechanisms (eds AWD Larkum, AR Grossman, JA Raven), pp. 27–55. Cham: Springer International Publishing.

15. Takizawa K, Kanazawa A, Cruz JA, Kramer DM. 2007 In vivo thylakoid proton motive force. Quantitative non-invasive probes show the relative lumen pH-induced regulatory responses of antenna and electron transfer. Biochim. Biophys. Acta 1767, 1233–1244.

16. Avenson TJ, Kanazawa A, Cruz JA, Takizawa K, Ettinger WE, Kramer DM. 2005 Integrating the proton circuit into photosynthesis: Progress and challenges. Plant Cell Environ. 28, 97–109.

17. Vershubskii AV, Tikhonov AN. 2020 pH-Dependent Regulation of Electron and Proton Transport in Chloroplasts In Situ and In Silico. Biochemistry (Moscow), Supplement Series A: Membrane and Cell Biology 14, 154–165.

18. Kanazawa A, Neofotis P, Davis GA, Fisher N, Kramer DM. 2020 Diversity in Photoprotection and Energy Balancing in Terrestrial and Aquatic Phototrophs. In Photosynthesis in Algae: Biochemical and Physiological Mechanisms (eds AWD Larkum, AR Grossman, JA Raven), pp. 299–327. Cham: Springer International Publishing.

19. Takahashi S, Milward SE, Fan DY, Chow WS, Badger MR. 2009 How does cyclic electron flow alleviate photoinhibition in Arabidopsis? Plant Physiol. 149, 1560–1567.

20. Chow WS, Hope AB. 1977 Proton Translocation, Electron Transport and Photophosphorylation in Isolated Chloroplasts. Plant Physiol. 4, 647–665.

21. Allahverdiyeva Y, Suorsa M, Tikkanen M, Aro EM. 2015 Photoprotection of photosystems in fluctuating light intensities. J. Exp. Bot. 66, 2427–2436.

22. Suorsa M et al. 2012 PROTON GRADIENT REGULATION5 Is Essential for Proper Acclimation of Arabidopsis Photosystem I to Naturally and Artificially Fluctuating Light Conditions. Plant Cell 24, 2934–2948.

23. Kanazawa A et al. 2017 Chloroplast ATP synthase modulation of the thylakoid proton motive force: Implications for photosystem I and photosystem II photoprotection. Frontiers in Plant Physiology https://doi.org/10.3389/fpls.2017.00719.

24. Chaux F, Burlacot A, Mekhalfi M, Auroy P, Blangy S, Richaud P, Peltier G. 2017 Flavodiiron Proteins Promote Fast and Transient O2 Photoreduction in Chlamydomonas. Plant Physiol. 174, 1825–1836.

25. Armbruster U, Leonelli L, Correa Galvis V, Strand D, Quinn EH, Jonikas MC, Niyogi KK. 2016 Regulation and Levels of the Thylakoid K+/H+ Antiporter KEA3 Shape the Dynamic Response of Photosynthesis in Fluctuating Light. Plant Cell Physiol. (doi:10.1093/pcp/pcw085)

26. Kulheim C, Agren J, Jansson S. 2002 Rapid regulation of light harvesting and plant fitness in the field. Science 297, 91–93.

27. Allahverdiyeva Y, Mustila H, Ermakova M, Bersanini L, Richaud P, Ajlani G, Battchikova N, Cournac L, Aro EM. 2013 Flavodiiron proteins Flv1 and Flv3 enable cyanobacterial growth and photosynthesis under fluctuating light. Proc. Natl. Acad. Sci. U. S. A. 110, 4111–4116.

28. Cruz JA, Savage LJ, Zegarac R, Hall CC, Satoh-Cruz M, Davis GA, Kovac WK, Chen J, Kramer DM. 2016 Dynamic environmental photosynthetic imaging reveals emergent phenotypes. Cell Syst 2, 365–377.

29. Davis GA et al. 2016 Limitations to photosynthesis by proton motive force-induced photosystem II photodamage. Elife eLife 2016;5:e16921.

30. Murchie EH, Niyogi KK. 2011 Manipulation of photoprotection to improve plant photosynthesis. Plant Physiol. 155, 86–92.

31. Gjindali A, Herrmann HA, Schwartz J-M, Johnson GN, Calzadilla PI. 2021 A Holistic Approach to Study Photosynthetic Acclimation Responses of Plants to Fluctuating Light. Front. Plant Sci. 12, 668512.

32. Zhu XG, Long SP, Ort DR. 2010 Improving photosynthetic efficiency for greater yield. Annu. Rev. Plant Biol. 61, 235–261.

33. Kuhlgert S et al. 2016 MultispeQ Beta: a tool for large-scale plant phenotyping connected to the open PhotosynQ network. R Soc Open Sci 3, 160592.

34. Loriaux SD, Avenson TJ, Welles JM, McDermitt DK, Eckles RD, Riensche B, Genty B. 2013 Closing in on maximum yield of chlorophyll fluorescence using a single multiphase flash of sub-saturating intensity. Plant Cell Environ. 2013/04/17. (doi:10.1111/pce.12115)

35. Genty B, Briantais J-M, Baker NR. 1989 The relationship between the quantum yield of photosynthetic electron transport and quenching of chlorophyll fluorescence. Biochim. Biophys. Acta 990, 87–92.

36. Kramer DM, Johnson G, Kiirats O, Edwards GE. 2004 New fluorescence parameters for the determination of Q_A_ redox state and excitation energy fluxes. Photosynth. Res. 79, 209–218.

37. Tietz S, Hall CC, Cruz JA, Kramer DM. 2017 NPQ(T): a chlorophyll fluorescence parameter for rapid estimation and imaging of non-photochemical quenching of excitons in photosystem II associated antenna complexes. Plant Cell Environ. 40, 1243–1255.

38. Kanazawa A, Kramer DM. 2002 In vivo modulation of nonphotochemical exciton quenching (NPQ) by regulation of the chloroplast ATP synthase. Proc. Natl. Acad. Sci. U. S. A. 99, 12789–12794.

39. Baker N, Harbinson J, Kramer DM. 2007 Determining the limitations and regulation of photosynthetic energy transduction in leaves. Plant Cell Environ. 30, 1107–1125.

40. Scrucca L, Fop M, Murphy TB, Raftery AE. 2016 mclust 5: clustering, classification and density estimation using Gaussian finite mixture models. R J.

41. Fraley C, Raftery AE. 2002 Model-Based Clustering, Discriminant Analysis, and Density Estimation. J. Am. Stat. Assoc. 97, 611–631.

42. Dasgupta A, Raftery AE. 1998 Detecting Features in Spatial Point Processes with Clutter via Model-Based Clustering. J. Am. Stat. Assoc. 93, 294–302.

43. Fraley C, Raftery AE. 1998 How Many Clusters? Which Clustering Method? Answers Via Model-Based Cluster Analysis. Comput. J. 41, 578–588.

44. Lambrev PH, Miloslavina Y, Jahns P, Holzwarth AR. 2012 On the relationship between non-photochemical quenching and photoprotection of Photosystem II. Biochim. Biophys. Acta 1817, 760–769.

45. Holzwarth AR, Miloslavina Y, Nilkens M, Jahns P. 2009 Identification of two quenching sites active in the regulation of photosynthetic light-harvesting studied by time-resolved fluorescence. Chem. Phys. Lett. 483, 262–267.

46. Davis GA et al. 2016 Limitations to photosynthesis by proton motive force-induced photosystem II photodamage. Elife 5. (doi:10.7554/eLife.16921)

47. Strand DD, Kramer •. D. M. 2014 Control of non-photochemical exciton quenching by the proton circuit of photosynthesis. In Non-Photochemical Quenching and Energy Dissipation in Plants, Algae and Cyanobacteria (eds B Demmig-Adams, G Garab, W Adams III, Govindjee), pp. 387–408. The Netherlands: Springer.

48. Foyer C, Furbank R, Harbinson J, Horton P. 1990 The mechanisms contributing to photosynthetic control of electron transport by carbon assimilation in leaves. Photosynth. Res. 25, 83–100.

49. Noctor G, Foyer CH. 2000 Homeostasis of adenylate status during photosynthesis in a fluctuating environment. J. Exp. Bot. 51, 347–356.

50. Stitt M. 1996 Metabolic regulation of photosynthesis. In Photosynthesis and the Environment (ed NR Baker), pp. 151–190. Dordrecht: Kluwer Academic Publishers.

51. Preiser AL, Fisher N, Banerjee A, Sharkey TD. 2019 Plastidic glucose-6-phosphate dehydrogenases are regulated to maintain activity in the light. Biochem. J 476, 1539–1551.

52. Cejudo FJ, Ojeda V, Delgado-Requerey V, González M, Pérez-Ruiz JM. 2019 Chloroplast Redox Regulatory Mechanisms in Plant Adaptation to Light and Darkness. Front. Plant Sci. 10, 380.

53. Hochmal AK, Schulze S, Trompelt K, Hippler M. 2015 Calcium-dependent regulation of photosynthesis. Biochim. Biophys. Acta 1847, 993–1003.

54. Takizawa K, Cruz JA, Kanazawa A, Kramer DM. 2007 The thylakoid proton motive force in vivo. Quantitative, non-invasive probes, energetics, and regulatory consequences of light-induced pmf. Biochim. Biophys. Acta 1767, 1233–1244.

55. Sonoike K, Terashima I. 1994 Mechanism of photosystem-I photoinhibition in leaves of Cucumis sativus L. Planta 194, 287–293.

56. Huang W, Sun H, Tan S-L, Zhang S-B. 2021 The water-water cycle is not a major alternative sink in fluctuating light at chilling temperature. Plant Sci. 305, 110828.

57. Lambrev PH, Tsonev T, Velikova V, Georgieva K, Lambreva MD, Yordanov I, Kovács L, Garab G. 2007 Trapping of the quenched conformation associated with non-photochemical quenching of chlorophyll fluorescence at low temperature. Photosynth. Res. 94, 321–332.

58. Demmig B. 1987 Photoinhibition and zeaxanthin formation in intact leaves. Plant Physiol. 84, 218–224.

59. Greer DH, Berry JA, Björkman O. 1986 Photoinhibition of photosynthesis in intact bean leaves: role of light and temperature, and requirement for chloroplast-protein synthesis during recovery. Planta 168, 253–260.

60. Bilger W, Björkman O. 1991 Temperature dependence of violaxanthin de-epoxidation and non-photochemical fluorescence quenching in intact leaves of Gossypium hirsutum L. and Malva parviflora L. Planta 184, 226–234.

61. Adams WW, Demmig-Adams B. 1995 The xanthophyll cycle and sustained thermal energy dissipation activity in Vinca minor and Euonymus kiautschovicus in winter. Plant, Cell and Environment. 18, 117–127. (doi:10.1111/j.1365-3040.1995.tb00345.x)

62. Verhoeven AS, Adams WW III, Demmig-Adams B. 1996 Close relationship between the state of the xanthophyll cycle pigments and photosystem II efficiency during recovery from winter stress. Physiologia Plantarum. 96, 567–576. (doi:10.1034/j.1399-3054.1996.960404.x)

63. Mullineaux CW, Pascal AA, Horton P, Holzwarth AR. 1993 Excitation-energy quenching in aggregates of the LHCII Chlorophyll-protein complex - a time-resolved fluorescence study. Biochim. Biophys. Acta 1141, 23–28.

64. Rees S, Young A, Noctor G, Horton P. 1989 Enhancement of the DpH-dependent dissipation of excitation energy in spinach chloroplasts by light-activation: correlation with the synthesis of zeaxanthin. FEBS Lett. 256, 85–90.

65. Horton P, Wentworth M, Ruban A. 2005 Control of the light harvesting function of chloroplast membranes: the LHCII-aggregation model for non-photochemical quenching. FEBS Lett. 579, 4201–4206.

66. Kirchhoff H, Hall C, Wood M, Herbstova M, Tsabari O, Nevo R, Charuvi D, Shimoni E, Reich Z. 2011 Dynamic control of protein diffusion within the granal thylakoid lumen. Proc. Natl. Acad. Sci. U. S. A. 108, 20248–20253.

67. Kaiser E, Morales A, Harbinson J. 2018 Fluctuating Light Takes Crop Photosynthesis on a Rollercoaster Ride. Plant Physiol. 176, 977–989.

68. Anderson CM et al. 2021 High Light and High Temperature Reduce Photosynthesis via Different Mechanisms in the C4 Model Setaria viridis. bioRxiv., 2021.02.20.431694. (doi:10.1101/2021.02.20.431694)

69. Stitt M, Zhu X-G. 2014 The large pools of metabolites involved in intercellular metabolite shuttles in C4 photosynthesis provide enormous flexibility and robustness in a fluctuating light environment. Plant Cell Environ. 37, 1985–1988.

70. Pearcy RW, Krall JP, Sassenrath-Cole GF. 1996 Photosynthesis in Fluctuating Light Environments. In Photosynthesis and the Environment (ed NR Baker), pp. 321–346. Dordrecht: Springer Netherlands.

71. Heineke D, Stitt M, Heldt HW. 1989 Effects of inorganic phosphate on the light dependent thylakoid energization of intact spinach chloropalsts. Plant Physiol., 221–226.

72. Thormählen I et al. 2017 Thioredoxins Play a Crucial Role in Dynamic Acclimation of Photosynthesis in Fluctuating Light. Mol. Plant 10, 168–182.

73. Carrillo LR, Froehlich JE, Cruz JA, Savage LJ, Kramer DM. 2016 Multi-level regulation of the chloroplast ATP synthase: The chloroplast NADPH thioredoxin reductase C (NTRC) is required for redox modulation specifically under low irradiance. Plant J. 87, 654–663.

74. Armbruster U, Correa Galvis V, Kunz H-H, Strand DD. 2017 The regulation of the chloroplast proton motive force plays a key role for photosynthesis in fluctuating light. Curr. Opin. Plant Biol. 37, 56–62.

75. Yin L, Lundin B, Bertrand M, Nurmi M, Solymosi K, Kangasjarvi S, Aro EM, Schoefs B, Spetea C. 2010 Role of thylakoid ATP/ADP carrier in photoinhibition and photoprotection of photosystem II in Arabidopsis. Plant Physiol. 153, 666–677.

76. Flood PJ et al. 2020 Reciprocal cybrids reveal how organellar genomes affect plant phenotypes. Nat Plants 6, 13–21.

