## Supplemental Data for "Light Potentials of Photosynthetic Energy Storage in the Field: What limits the ability to use or dissipate rapidly increased light energy?"

**Table S1A**

| Covariates | P value in explaining $LEF_{amb}$ |
|---|---|
| $\sqrt{PAR}$ | <2e-16 *** |
| $T_{leaf}$ | <2e-16 *** |
| $\sqrt{PAR}$ : $T_{leaf}$ | <2e-16 *** |

| Covariates | P value in explaining $LEF_{high}$ |
|---|---|
| $\sqrt{PAR}$ | <2e-16 *** |
| $T_{leaf}$ | <2e-16 *** |
| $\sqrt{PAR}$ : $T_{leaf}$ | 1.41 e-05 *** |

**Table S1B**

| Covariates | P value in explaining $NPQ_{amb}$ |
|---|---|
| $\sqrt{PAR}$ | 0.000473 *** |
| $T_{leaf}$ | 0.912335 |
| $\sqrt{PAR}$ : $T_{leaf}$ | 1.17e-06 *** |

| Covariates | P value in explaining $NPQ_{high}$ |
|---|---|
| $\sqrt{PAR}$ | 2.43e-06 *** |
| $T_{leaf}$ | <2e-16 *** |
| $\sqrt{PAR}$ : $T_{leaf}$ | 0.0262 * |

**Tables S1A and B.** Simple Linear Models to express the association between a photosynthetic trait and model covariates. Column1 lists the covariates used in the model: Square root values of $PAR_{amb}$ ($\sqrt{PAR}$), leaf temperature ($T_{Leaf}$) and interaction between $PAR_{amb}$ and $T_{Leaf.}$ Column 2 expresses p-values related with each covariate. *** represents covariates being highly significant, * representing covariates being significant at 5% level.

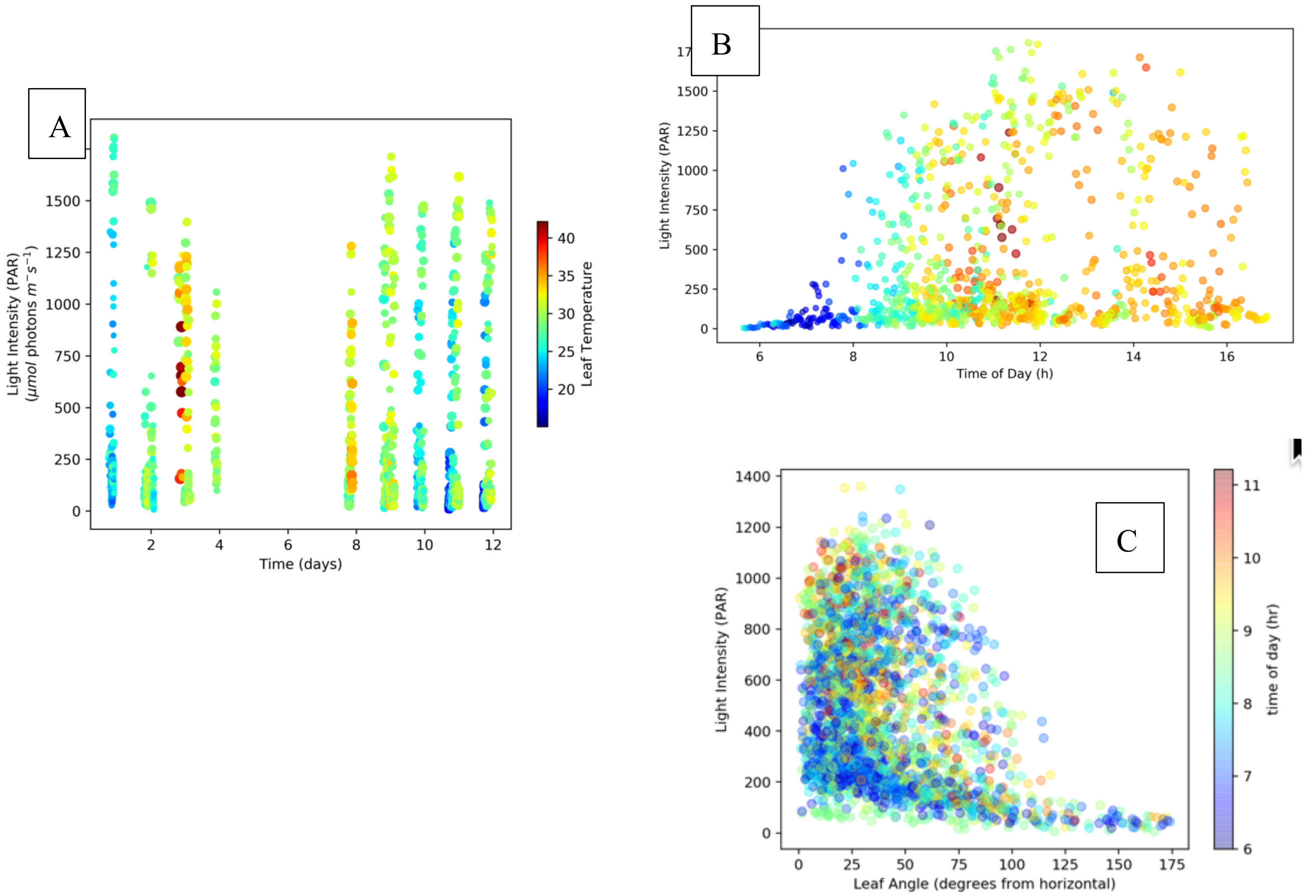


**Figure S1**. Dependencies of ambient Photosynthetically active radiation (Light intensity, $PAR_{amb}$) and leaf temperature ($T_{leaf}$) plotted over time over the course of the entire experiment (Panel A) or aggregated into a single apparent day (Panel B). Panel C plots $PAR_{amb}$ as functions of both leaf angle in degrees from parallel to the earth surface (X-axis) and time of day (symbol color).

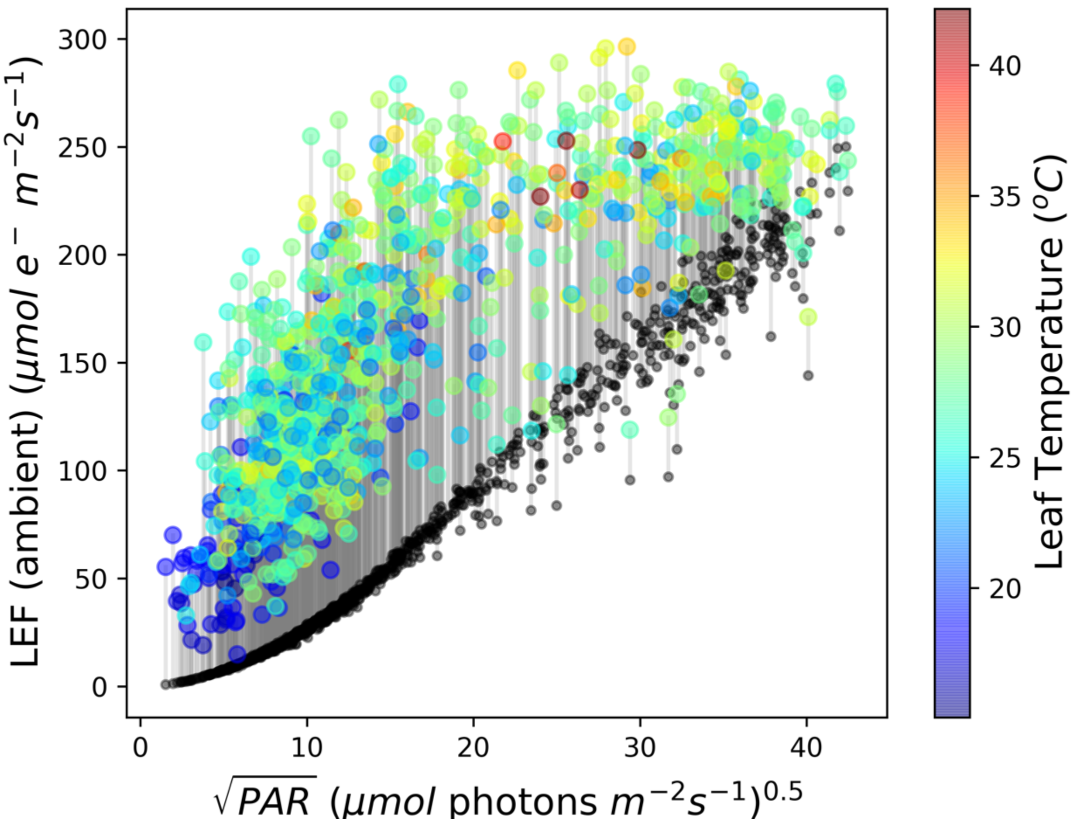


**Figure S2**. The dependence of LEF under ambient PAR ($LEF_{amb}$) and after 10s at 2000 μmol photons $m^{-2}$ $s^{-1}$ ($PAR_{high}$) as functions of the square root of $PAR_{amb}$ (X-axis) and leaf temperature ($T_{leaf}$) (see coloration).

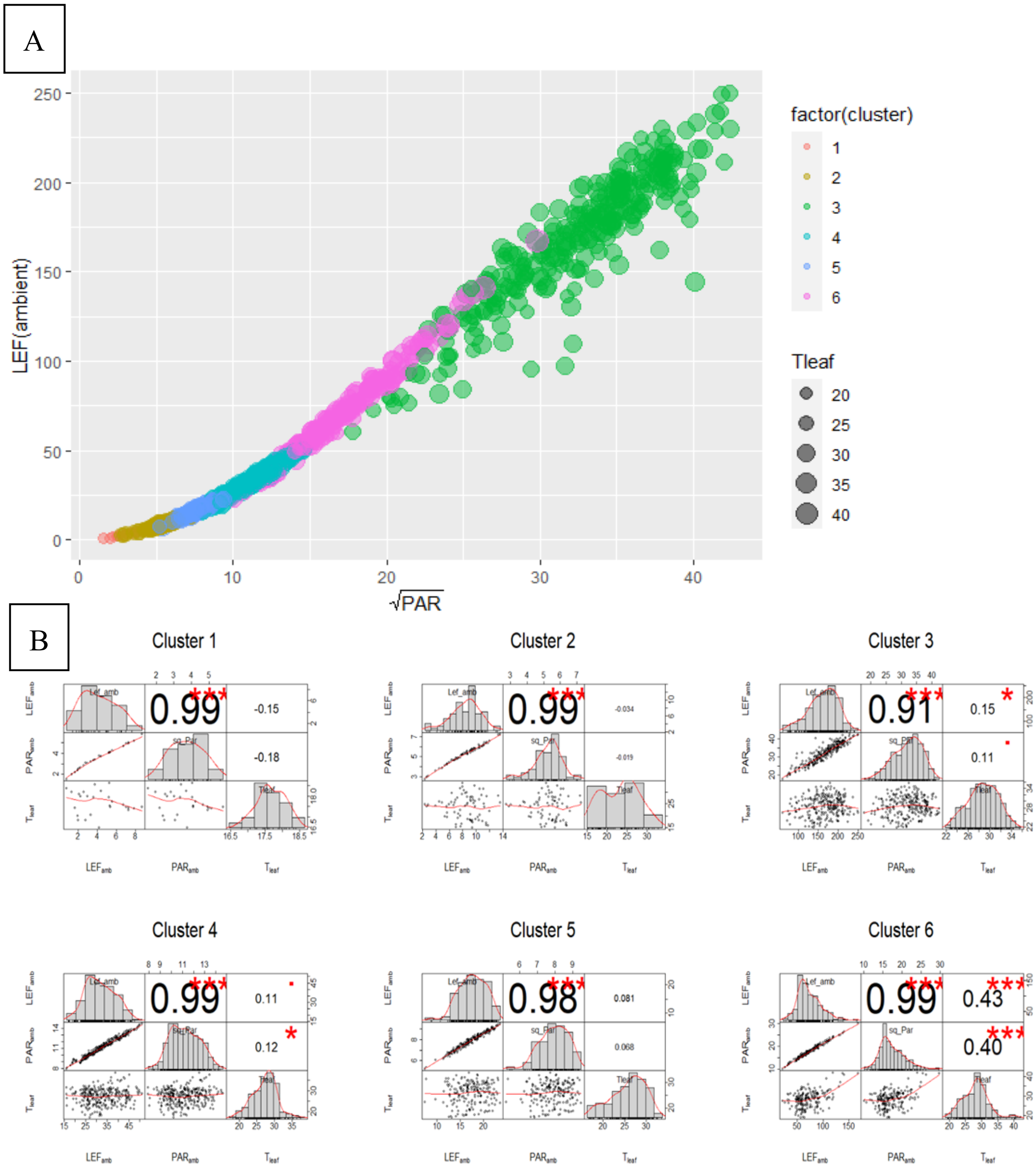


**Figure S3.** Gaussian Mixture Model (GMM) clustering of $LEF_{amb}$ (Panel A) and correlation matrixes between $LEF_{amb}$, $PAR_{amb}$ and leaf temperature ($T_{leaf}$) for each cluster (Panel B). Panel A contains the clustering plot for specific phenotypes with environmental parameters ($PAR_{amb}$ and $T_{Leaf}$). The different subplots in Panel B represent the co-dependence between the parameters used. For each subplot the diagonal entries are the density plots for individual parameters, lower triangular entries represent the plots taking two parameters at a time and upper triangular entries are the Pearson correlation coefficients for the selected parameters. Significance is indicated by $p<0.001$ (***), $0.001<p<0.01$ (**), $0.01<p<0.05$ (*), $0.05<p<0.1$ (.) and $p>0.1$ (no symbol).

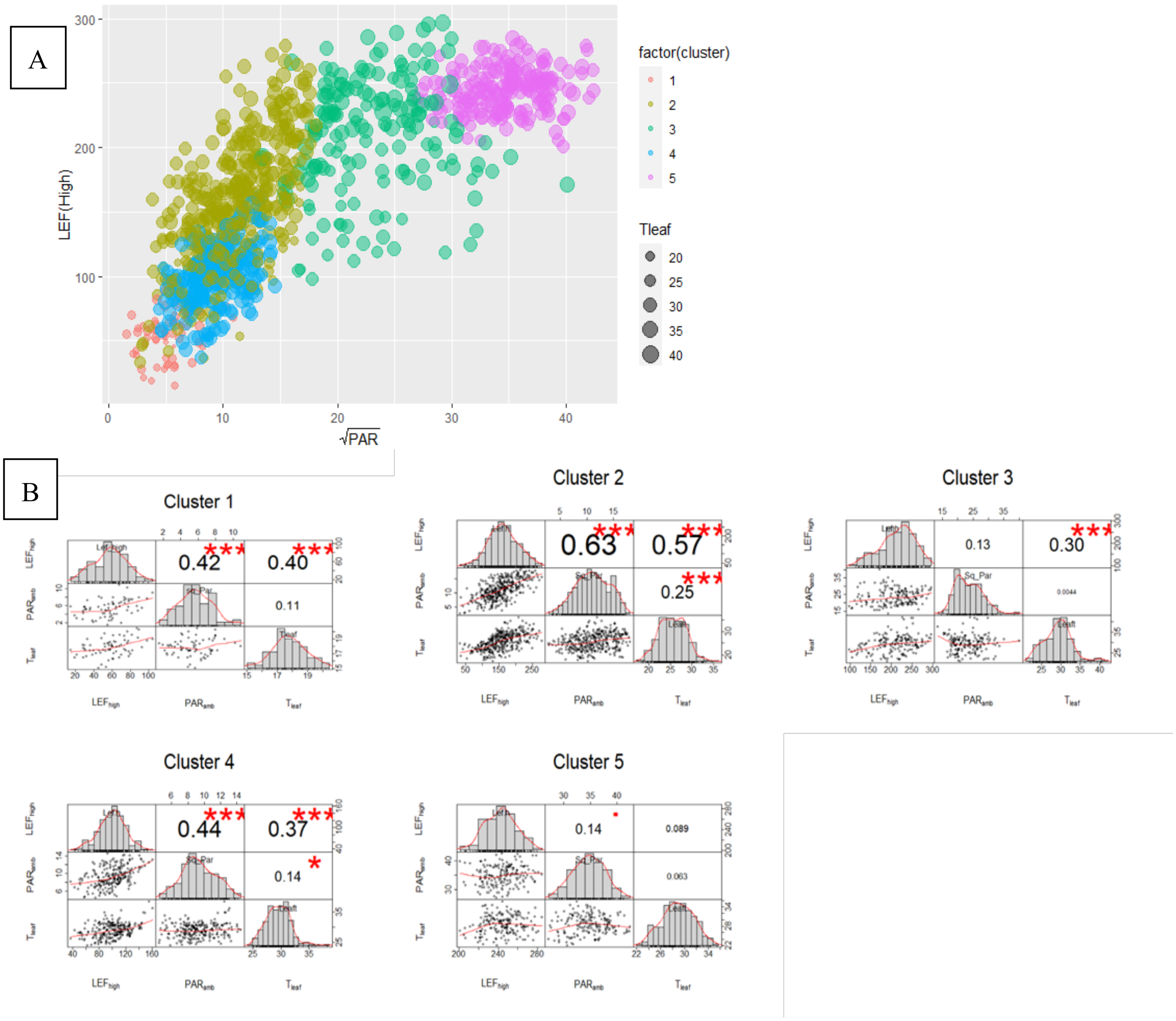


**Figure S4.** Gaussian Mixture Model (GMM) clustering of $LEF_{High}$ (Panel A) and correlation matrixes between $LEF_{High}$, $PAR_{amb}$ and leaf temperature ($T_{leaf}$) for each cluster (Panel B). Panel A contains the clustering plot for specific phenotypes with environmental parameters ($PAR_{amb}$ and $T_{Leaf}$). The different subplots in Panel B represent the co-dependence between the parameters used. For each subplot the diagonal entries are the density plots for individual parameters, lower triangular entries represent the plots taking two parameters at a time and upper triangular entries are the Pearson correlation coefficients for the selected parameters. Significance is indicated by $p<0.001$ (***), $0.001<p<0.01$ (**), $0.01<p<0.05$ (*), $0.05<p< 0.1$ (.) and $p>0.1$ (no symbol).

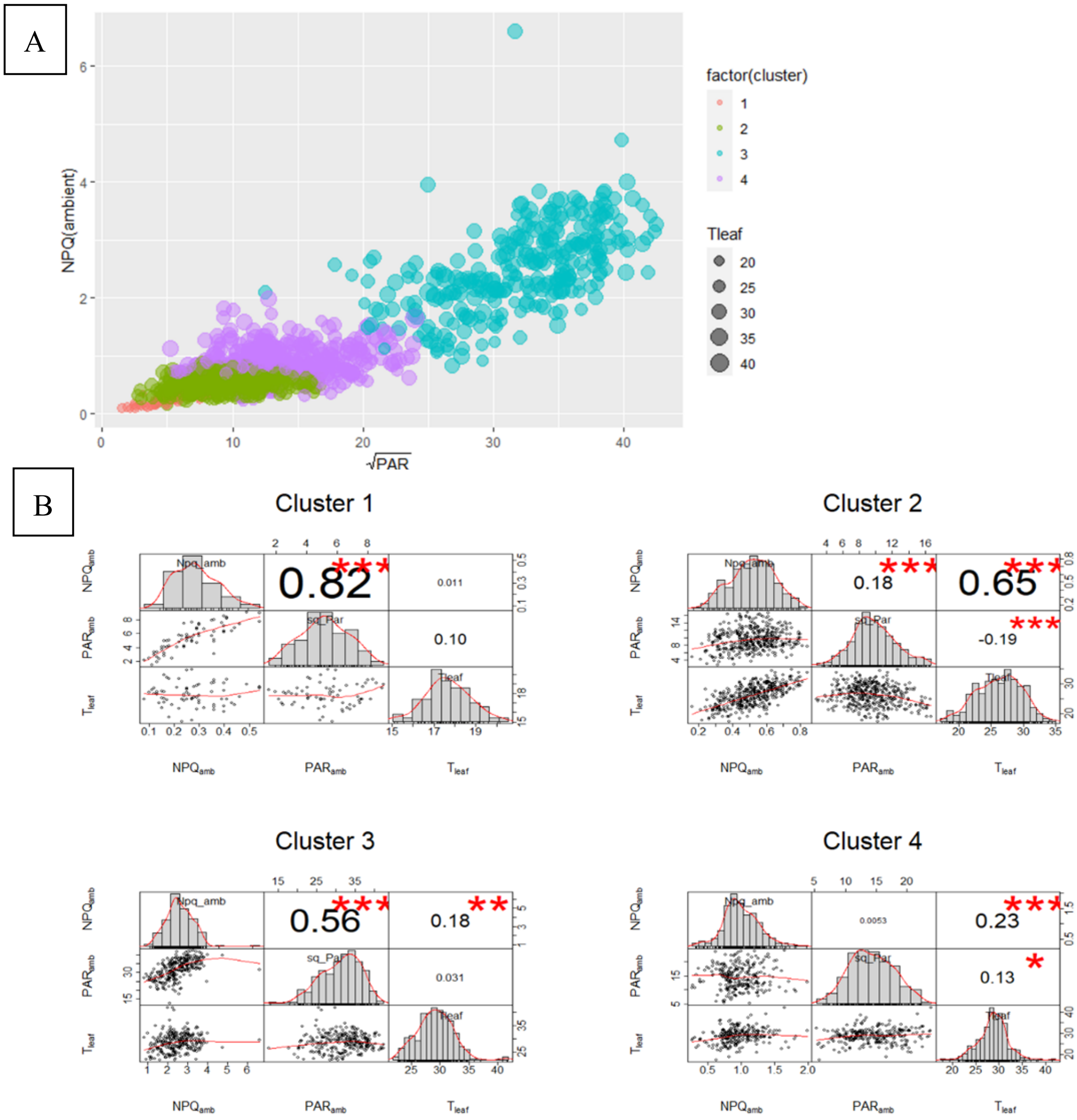


**Figure S5.** Gaussian Mixture Model (GMM) clustering of $NPQ_{amb}$ (Panel A) and correlation matrixes between $NPQ_{amb}$, $PAR_{amb}$ and leaf temperature ($T_{leaf}$) for each cluster (Panel B). Panel A contains the clustering plot for specific phenotypes with environmental parameters ($PAR_{amb}$ and $T_{Leaf}$). The different subplots in Panel B represent the co-dependence between the parameters used. For each subplot the diagonal entries are the density plots for individual parameters, lower triangular entries represent the plots taking two parameters at a time and upper triangular entries are the Pearson correlation coefficients for the selected parameters. Significance is indicated by $p<0.001$ (***), $0.001<p<0.01$ (**), $0.01<p<0.05$ (*), $0.05<p< 0.1$ (.) and $p>0.1$ (no symbol).

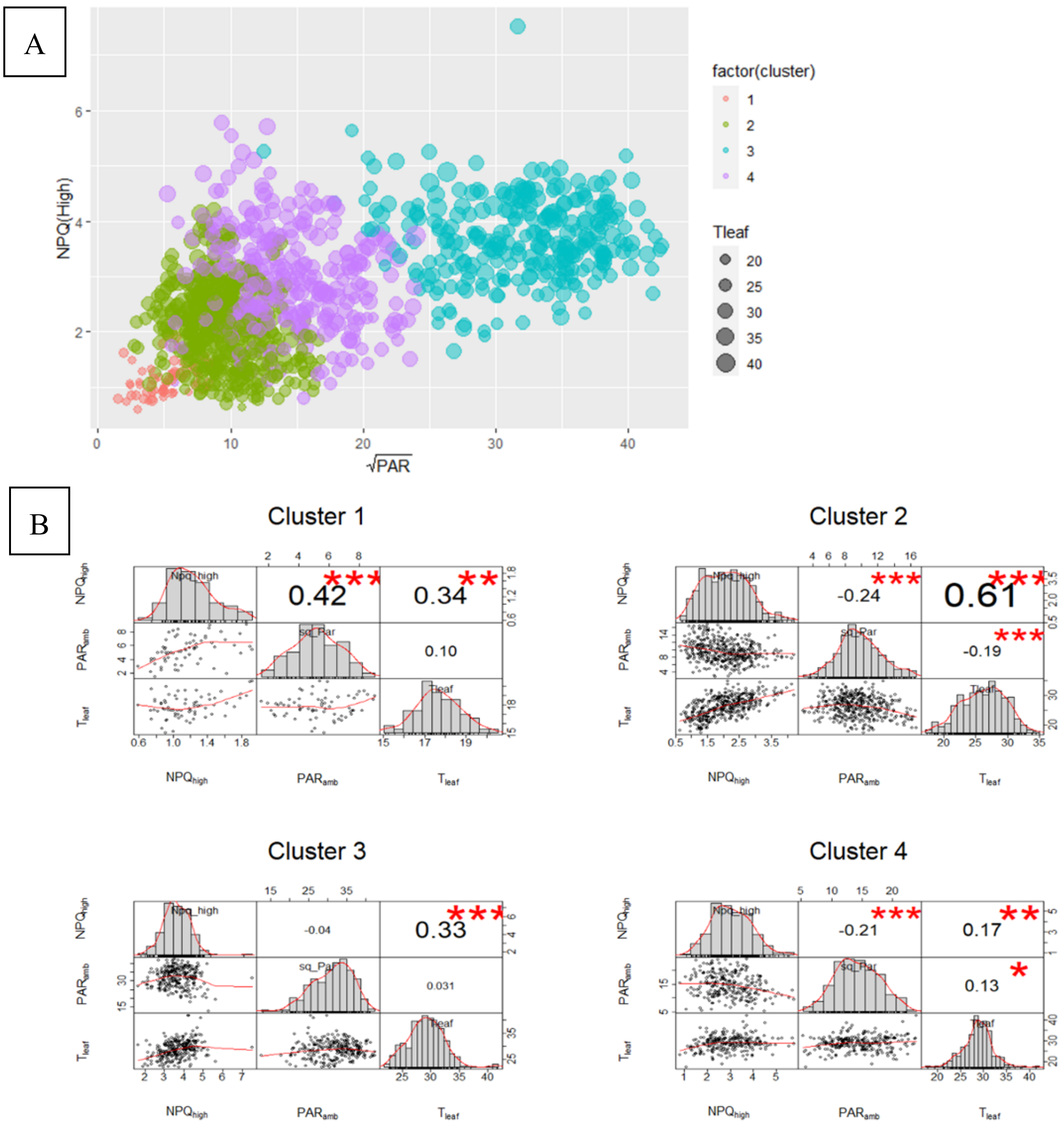


**Figure S6.** Gaussian Mixture Model (GMM) clustering of $NPQ_{High}$ (Panel A) and correlation matrixes between $NPQ_{High}$, $PAR_{amb}$ and leaf temperature ($T_{leaf}$) for each cluster (Panel B). Panel A contains the clustering plot for specific phenotypes with environmental parameters ($PAR_{amb}$ and $T_{Leaf}$). The different subplots in Panel B represent the co-dependence between the parameters used. For each subplot the diagonal entries are the density plots for individual parameters, lower triangular entries represent the plots taking two parameters at a time and upper triangular entries are the Pearson correlation coefficients for the selected parameters. Significance is indicated by $p<0.001$ (***), $0.001<p<0.01$ (**), $0.01<p<0.05$ (*), $0.05<p<0.1$ (.) and $p>0.1$ (no symbol).

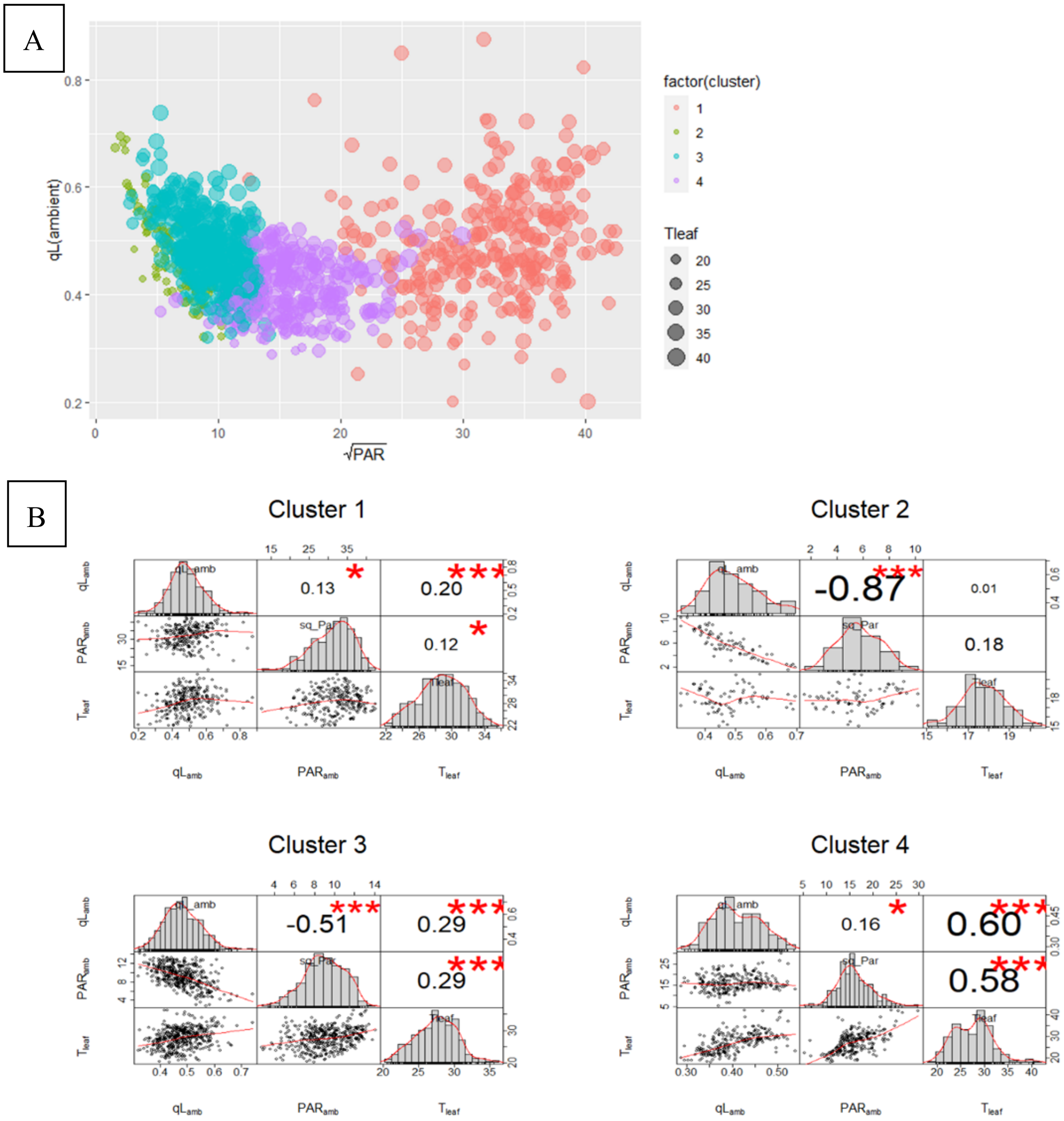


**Figure S7.** Gaussian Mixture Model (GMM) clustering of $q_{L(ambient)}$ (Panel A) and correlation matrixes between $q_{L(ambient)}$, $PAR_{amb}$ and leaf temperature ($T_{leaf}$) for each cluster (Panel B). Panel A contains the clustering plot for specific phenotypes with environmental parameters ($PAR_{amb}$ and $T_{Leaf}$). The different subplots in Panel B represent the co-dependence between the parameters used. For each subplot the diagonal entries are the density plots for individual parameters, lower triangular entries represent the plots taking two parameters at a time and upper triangular entries are the Pearson correlation coefficients for the selected parameters. Significance is indicated by $p<0.001$ (***), $0.001<p<0.01$ (**), $0.01<p<0.05$ (*), $0.05<p< 0.1$ (.) and $p>0.1$ (no symbol).

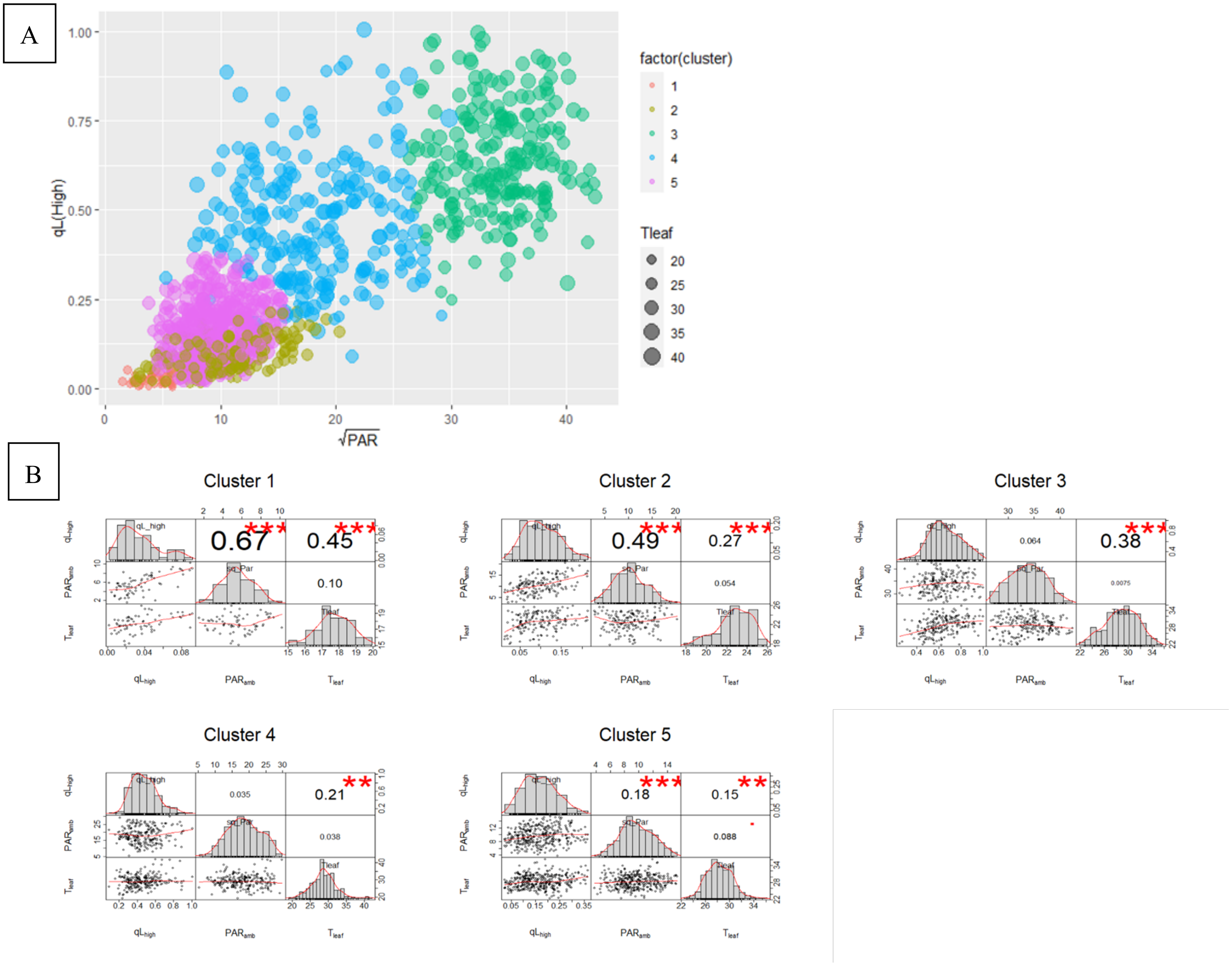


**Figure S8.** Gaussian Mixture Model (GMM) clustering of $q_{L(High)}$ (Panel A) and correlation matrixes between $q_{L(High)}$, $PAR_{amb}$ and leaf temperature ($T_{leaf}$) for each cluster (Panel B). Panel A contains the clustering plot for specific phenotypes with environmental parameters ($PAR_{amb}$ and $T_{Leaf}$). The different subplots in Panel B represent the co-dependence between the parameters used. For each subplot the diagonal entries are the density plots for individual parameters, lower triangular entries represent the plots taking two parameters at a time and upper triangular entries are the Pearson correlation coefficients for the selected parameters. Significance is indicated by $p<0.001$ (***), $0.001<p<0.01$ (**), $0.01<p<0.05$ (*), $0.05<p< 0.1$ (.) and $p>0.1$ (no symbol).

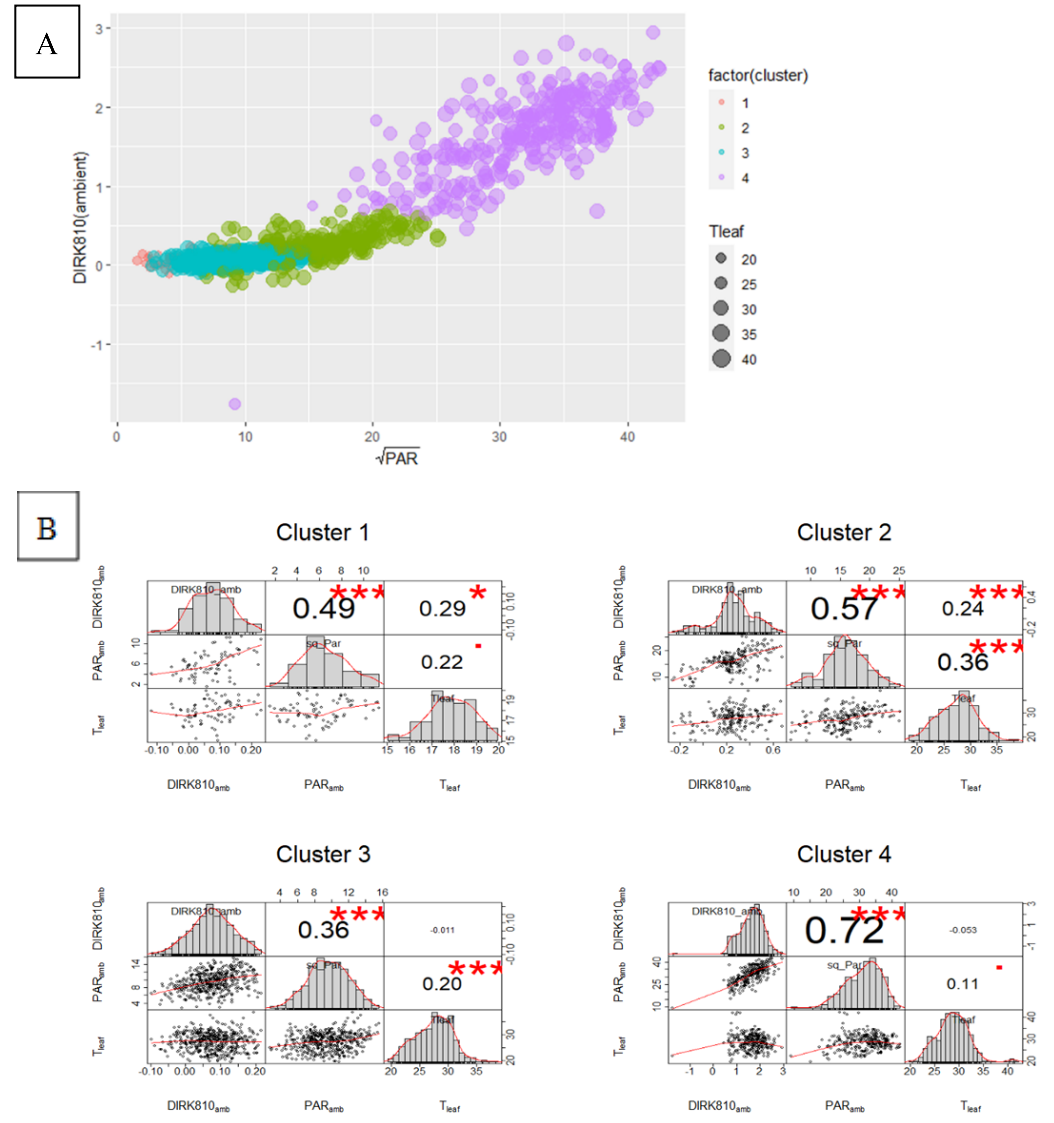


**Figure S9.** Gaussian Mixture Model (GMM) clustering of $P_{700}+_{(amb)}$ (here in the plot represented as $DIRK810_{amb}$) (Panel A) and correlation matrixes between $P_{700}+_{(ambient)}$, $PAR_{amb}$ and leaf temperature ($T_{leaf}$) for each cluster (Panel B). Panel A contains the clustering plot for specific phenotypes with environmental parameters ($PAR_{amb}$ and $T_{Leaf}$). The different subplots in Panel B represent the co-dependence between the parameters used. For each subplot the diagonal entries are the density plots for individual parameters, lower triangular entries represent the plots taking two parameters at a time and upper triangular entries are the Pearson correlation coefficients for the selected parameters. Significance is indicated by $p<0.001$ (***), $0.001<p<0.01$ (**), $0.01<p<0.05$ (*), $0.05<p< 0.1$ (.) and $p>0.1$ (no symbol).

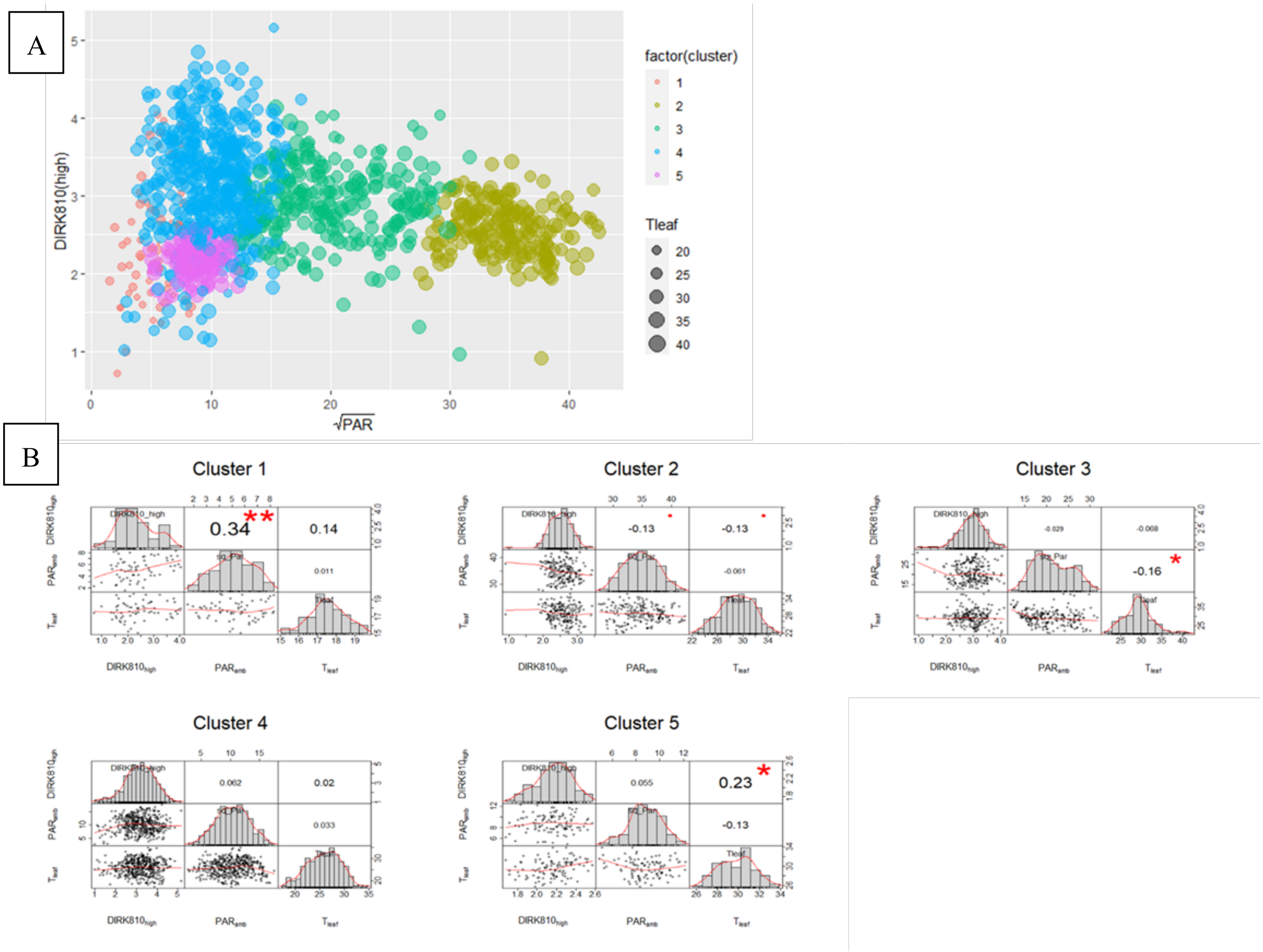


**Figure S10.** Gaussian Mixture Model (GMM) clustering of $P_{700}+_{(High)}$ (here in the plot represented as $DIRK810_{High}$ ) (Panel A) and correlation matrixes between $P_{700}+_{(high)}$, $PAR_{amb}$ and leaf temperature ($T_{leaf}$) for each cluster (Panel B). Panel A contains the clustering plot for specific phenotypes with environmental parameters ($PAR_{amb}$ and $T_{Leaf}$). The different subplots in Panel B represent the co-dependence between the parameters used. For each subplot the diagonal entries are the density plots for individual parameters, lower triangular entries represent the plots taking two parameters at a time and upper triangular entries are the Pearson correlation coefficients for the selected parameters. Significance is indicated by p<0.001 (***), 0.001<p<0.01 (**), 0.01<p<0.05 (*), 0.05<p< 0.1 (.) and p>0.1 (no symbol).

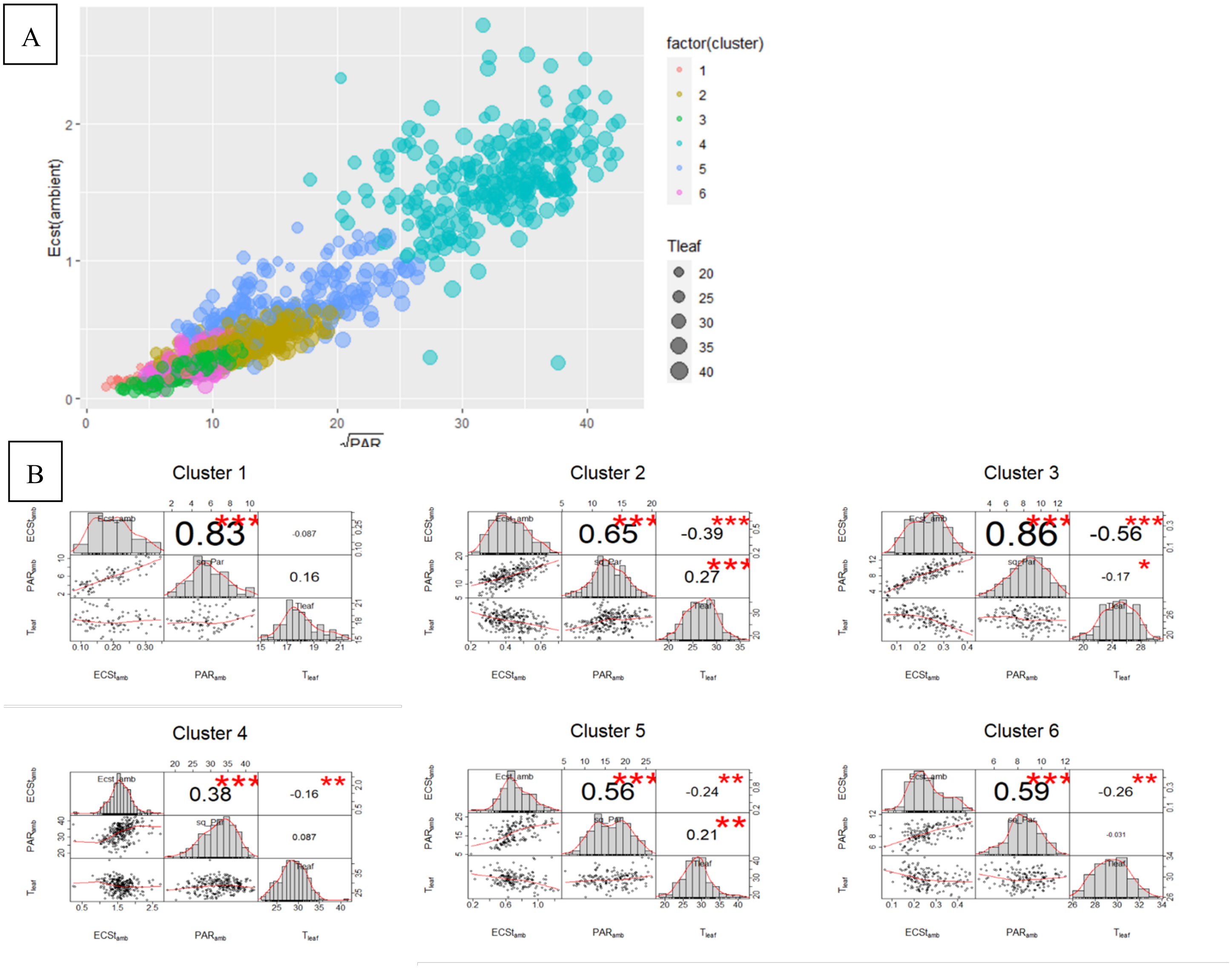


**Figure S11.** Gaussian Mixture Model (GMM) clustering of $ECS_{t(ambient)}$ (Panel A) and correlation matrixes between $ECS_{t(Ambient)}$, $PAR_{amb}$ and leaf temperature ($T_{leaf}$) for each cluster (Panel B). Panel A contains the clustering plot for specific phenotypes with environmental parameters ($PAR_{amb}$ and $T_{Leaf}$). The different subplots in Panel B represent the co-dependence between the parameters used. For each subplot the diagonal entries are the density plots for individual parameters, lower triangular entries represent the plots taking two parameters at a time and upper triangular entries are the Pearson correlation coefficients for the selected parameters. Significance is indicated by p<0.001 (***), 0.001<p<0.01 (**), 0.01<p<0.05 (*), 0.05<p< 0.1 (.) and p>0.1 (no symbol).

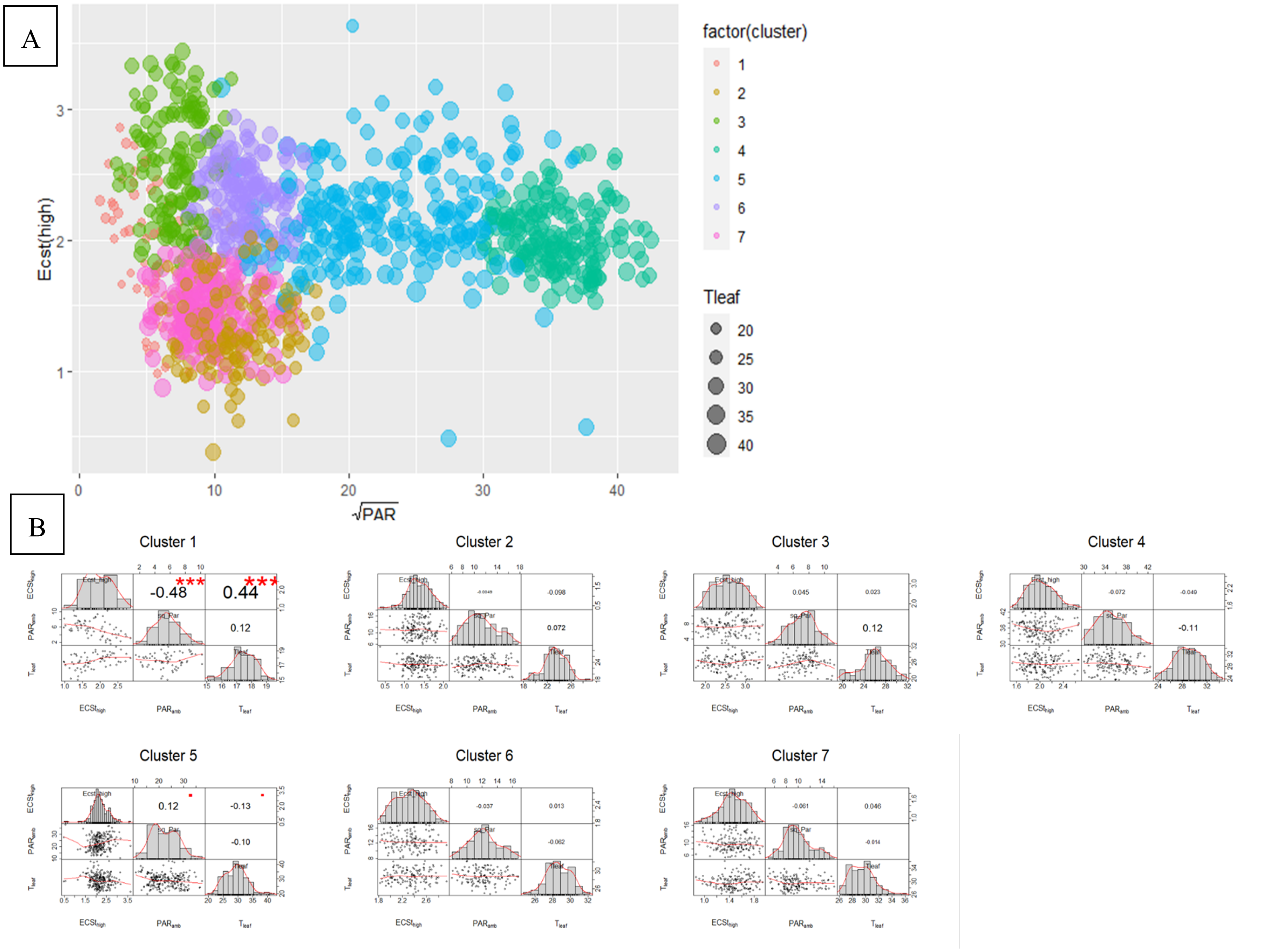


**Figure S12.** Gaussian Mixture Model (GMM) clustering of $ECS_{t(High)}$ (Panel A) and correlation matrixes between $ECS_{t(High)}$, $PAR_{amb}$ and leaf temperature ($T_{leaf}$) for each cluster (Panel B). Panel A contains the clustering plot for specific phenotypes with environmental parameters ($PAR_{amb}$ and $T_{Leaf}$). The different subplots in Panel B represent the co-dependence between the parameters used. For each subplot the diagonal entries are the density plots for individual parameters, lower triangular entries represent the plots taking two parameters at a time and upper triangular entries are the Pearson correlation coefficients for the selected parameters. Significance is indicated by $p<0.001$ (***), $0.001<p<0.01$ (**), $0.01<p<0.05$ (*), $0.05<p< 0.1$ (.) and $p>0.1$ (no symbol).

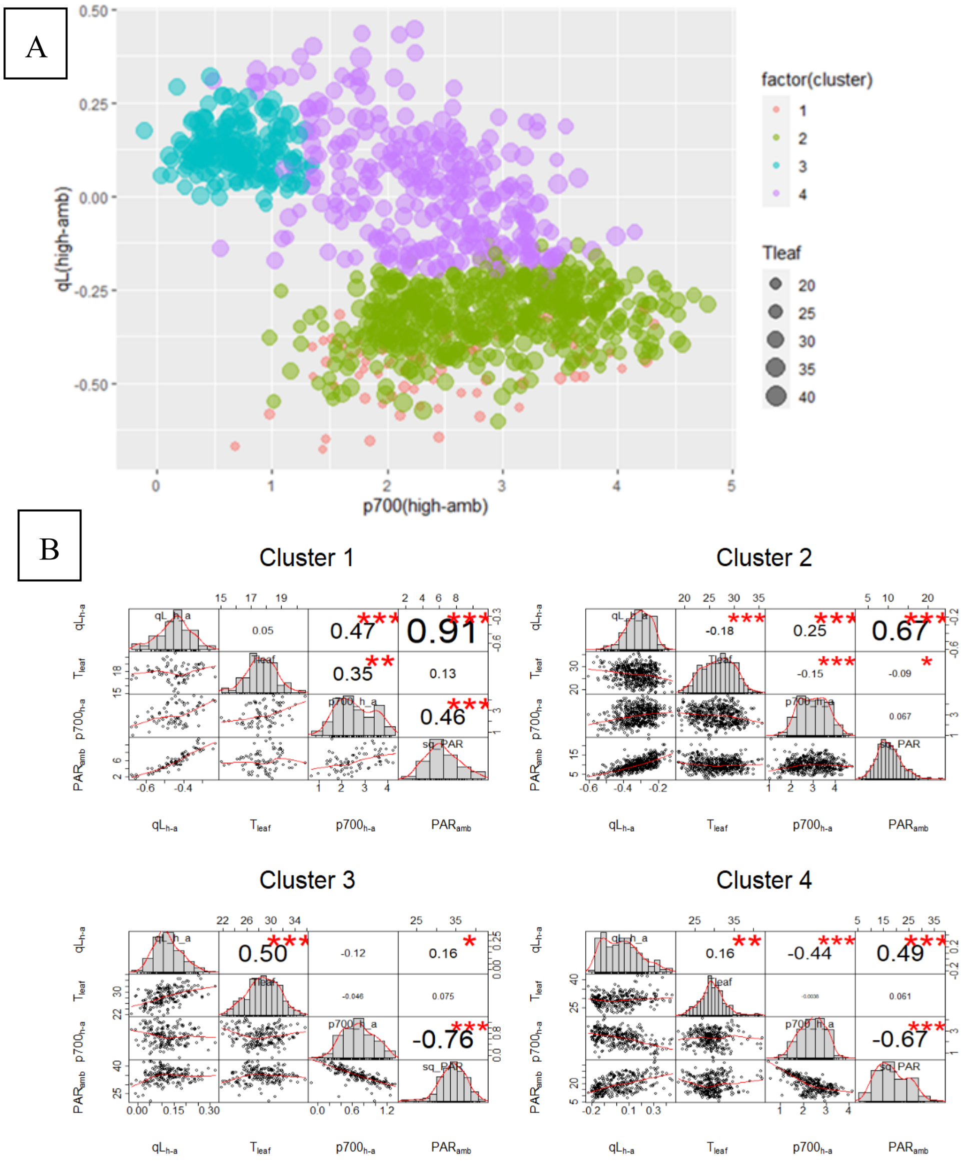


**Figure S13.** Gaussian Mixture Model (GMM) clustering with inputs as $qL_{(\text{high-ambient})}$ , $p700_{(\text{high-ambient})}$ and leaf temperature ($T_{leaf}$). (Panel A: plotted against adjusted $p700_{(\text{high-ambient})}$ values) and correlation matrixes between $qL_{(\text{high-ambient})}$, $PAR_{amb}$, leaf temperature ($T_{leaf}$), $p700_{(\text{high-ambient})}$ for each cluster (Panel B). Panel A contains the clustering plot for specific input parameters. The different subplots in Panel B represent the co-dependence between the parameters used. For each subplot the diagonal entries are the density plots for individual parameters, lower triangular entries represent the plots taking two parameters at a time and upper triangular entries are the Pearson correlation coefficients for the selected parameters. Significant level codes are given by: Significant is indicated by $p<0.0001$ (***), $0.0001<p<0.001$ (**), $0.001<p<0.01$ (*), $0.01<p< 0.05$ (.) and $p>0.05$ (no symbol).
